# Prefrontal cortex represents heuristics that shape choice bias and its integration into future behavior

**DOI:** 10.1101/2020.03.20.000224

**Authors:** Gabriela Mochol, Roozbeh Kiani, Rubén Moreno-Bote

**Affiliations:** Center for Brain and Cognition and Department of Information and Communications Technologies, Pompeu Fabra University, Barcelona, Spain; Center for Neural Science, New York University, New York, NY 10003, USA; Neuroscience Institute, NYU Langone Medical Center, New York, NY 10016, USA; Department of Psychology, New York University, New York, NY 10003, USA

**Keywords:** behavioral bias, decision making, prefrontal cortex, macaque monkey

## Abstract

Goal-directed behavior requires integrating sensory information with prior knowledge about the environment. Behavioral biases that arise from these priors could increase positive outcomes when the priors match the true structure of the environment, but mismatches also happen frequently and could cause unfavorable outcomes. Biases that reduce gains and fail to vanish with training indicate fundamental suboptimalities arising from ingrained heuristics of the brain. Here, we report systematic, gain-reducing choice biases in highly-trained monkeys performing a motion direction discrimination task where only the current stimulus is behaviorally relevant. The monkey’s bias fluctuated at two distinct time scales: slow, spanning tens to hundreds of trials, and fast, arising from choices and outcomes of the most recent trials. Our finding enabled single trial prediction of biases, which influenced the choice especially on trials with weak stimuli. The pre-stimulus activity of neuronal ensembles in the monkey prearcuate gyrus represented these biases as an offset along the decision axis in the state space. This offset persisted throughout the stimulus viewing period, when sensory information was integrated, leading to a biased choice. The pre-stimulus representation of history-dependent bias was functionally indistinguishable from the neural representation of upcoming choice before stimulus onset, validating our model of single-trial biases and suggesting that pre-stimulus representation of choice could be fully defined by biases inferred from behavioral history. Our results indicate that the prearcuate gyrus reflects intrinsic heuristics that compute bias signals, as well as the mechanisms that integrate them into the oculomotor decision-making process.

## Introduction

Choice biases are prevalent (Gardner, 2019). Biases that reflect imbalanced priors or reward expectations in the environment are advantageous as they could improve the speed or overall gain of our choices (Averbeck et al., 2006; Fan et al., 2018; Hanks et al., 2011; Hermoso-Mendizabal et al., 2020; Hesselmann et al., 2008; Moreno-Bote et al., 2008, 2011). However, biases could also hinder performance (Akrami et al., 2018), especially when they arise from heuristics that do not truly capture the environment or task structure. It is often hypothesized that the history of past stimuli, actions and outcomes inform these heuristics for sequential choices, where subjects perform a series of similar decisions (Abrahamyan et al., 2016; Akrami et al., 2018; Busse et al., 2011; Eskandar and Assad, 1999; Fritsche et al., 2017; Hermoso-Mendizabal et al., 2020; Hwang et al., 2017; Lueckmann et al., 2018; Nogueira et al., 2017; Padoa-Schioppa, 2013; Shadlen and Newsome, 2001; Wyart and Tallon-Baudry, 2009). However, the computations involved in these heuristics are not well defined. Further, the neural representation of history dependent biases and how they integrate in the decision-making process remain debated.

Addressing these questions requires developing a task in which heuristic biases are not rewarding. Biases that increase reward rate encourage alteration of decision strategies based on task structure, complicating generalization of results across tasks. But systematic biases that persist with training and are non-rewarding (or even reduce gain) provide an opportunity to explore the heuristics that shape history-dependent biases. Addressing our questions also requires single trial quantification of the magnitude of bias, as well as recording from neural ensembles in brain regions that represent both the bias and the decision-making process. Single trial quantification of the magnitude of bias necessitates development of behavioral models that can accurately predict the bias on individual trials. Although many studies attempted to do so (Gold et al., 2008a; Jasper et al., 2019; Lueckmann et al., 2018), there are few comprehensive models that achieve sufficient accuracy, often because they ignore one or more key factors that shape the bias. Additionally, past studies on the neural representation of bias focused largely on single neuron activity (Eskandar and Assad, 1999; Hanks et al., 2011; Lueckmann et al., 2018; Nogueira et al., 2017; Padoa-Schioppa, 2013; Shadlen and Newsome, 2001), whereas accurate characterization of the moment-by-moment fluctuation of the state of the neural population requires simultaneous recording of many neurons (Arandia-Romero et al., 2017; Jasper et al., 2019; Kiani et al., 2014a). Finally, past studies focused largely on finding a neural representation of bias and rarely explored how the bias is integrated in the decision-making process. Here, we overcome these challenges for the first time, by developing a comprehensive framework that determines the magnitude of bias on individual trials, characterizes bias representation by the prefrontal neural population, and determines the computational mechanism for integrating the bias in the decision-making process.

We recorded simultaneously from large populations of pre-arcuate gyrus neurons while monkeys performed a direction discrimination task, designed such that past history was unrelated to the present stimulus, making history-dependent biases suboptimal. Nonetheless and despite extensive training, monkeys showed detectable biases that fluctuated throughout and across experimental sessions. Their choice biases stemmed from two sources: fast biases shaped by actions and outcomes of past trials, especially the most recent one, and slow biases fluctuating over tens to hundreds of trials. Our single-trial quantification of bias enabled us to show that the total bias, as well as its two sources, were represented in the population activity of pre-arcuate gyrus neurons prior to the stimulus onset. The same pre-stimulus neural responses were also predictive of the upcoming choice. These neuronal representations of bias and choice were well aligned in the activity state space, suggesting that the pre-stimulus choice prediction was achieved through the representation of bias, which itself reflected past choices and feedbacks. Finally, we demonstrate that the bias influenced the decision-making process as an initial offset in the accumulation of evidence, pointing at the computational mechanism for the integration of bias in the decision-making process.

## Results

Using a 96-channel multi-electrode array, we recorded neural population activity from the prearcuate gyrus (area 8Ar, PAG), while monkeys (*n* = 2) performed a direction discrimination task (16 sessions; (Kiani et al., 2014b, 2015)). Each trial began with the monkey fixating on a central fixation point on the screen, following by the appearance of two targets (T1 and T2; Figure 1A), and a circular patch of random dot kinematogram (Britten et al., 1992). The percentage of coherently moving dots (coherence or motion strength) and the net motion direction of motion varied randomly trial to trial. The motion stimulus was shown for 800 ms and was followed by a delay period. The monkey reported motion direction at the end of the delay period with a saccadic eye movement to the corresponding targets. We use signed motion coherence (Britten et al., 1992; Kiani et al., 2008; Shadlen and Newsome, 2001) to jointly represent the stimulus strength and direction with a single variable (positive for motion toward T1 and negative for motion toward T2).

**Figure 1.**
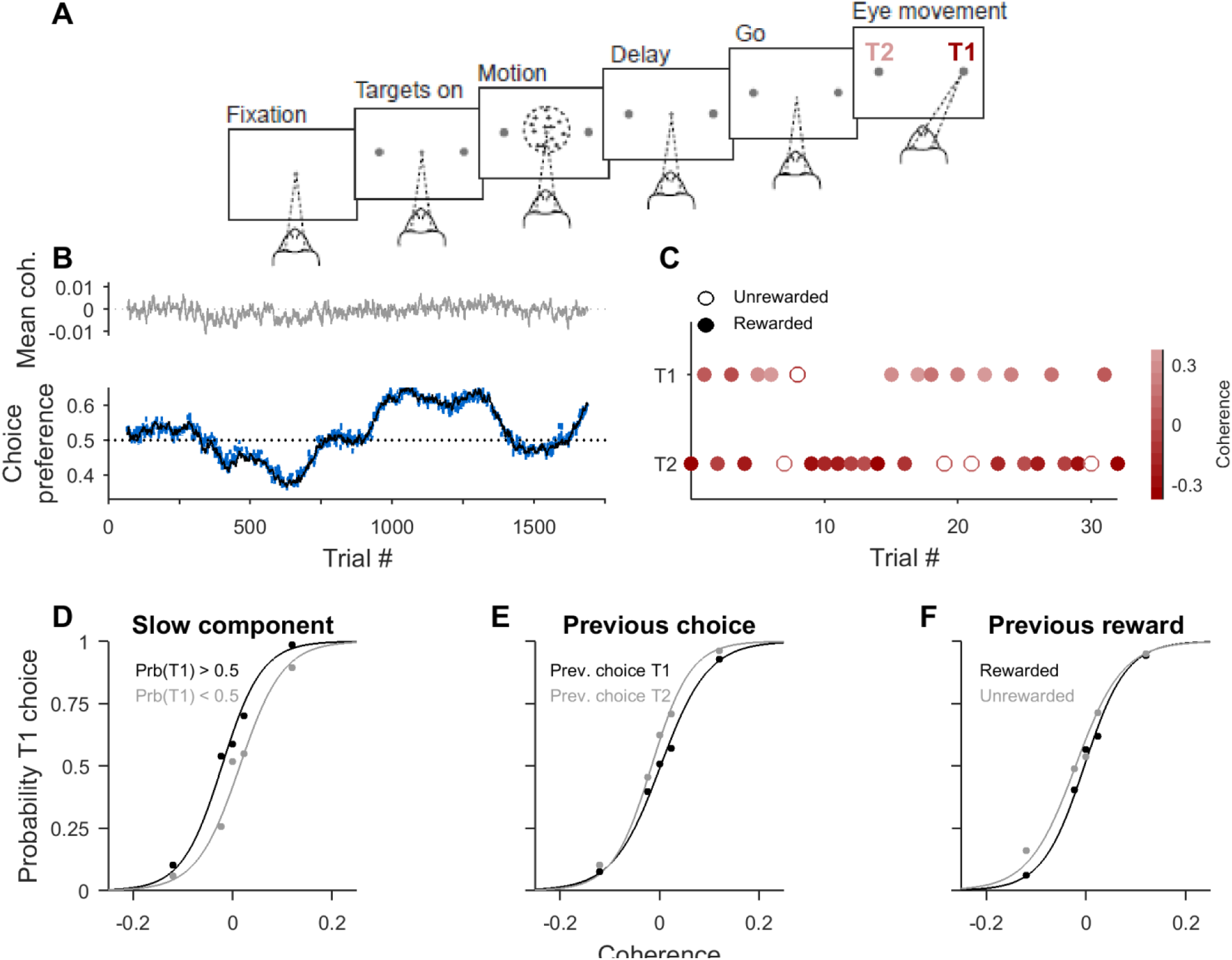
Well-trained monkeys in a random dots direction-discrimination task show slow and fast fluctuations in their preferred choice. **A**. Task design. After the monkey acquired a central fixation point, the patch of randomly moving dots appeared on the screen for 800 ms. The fraction of dots moving coherently in a given direction defined the trial difficulty. The motion was followed by a delay period with variable length, after which the monkey indicated its choice by making a saccade towards one of the two targets (T1 or T2). **B**. Top –average of signed motion coherence. Dashed line indicates 0% coherence, and positive and negative values indicate motion toward T1 and T2, respectively. Bottom – black line shows the fraction of trials in which the monkey chose T1. The dashed blue line also shows the choice preference towards T1 but calculated after balancing coherence in a given window. All curves calculated in a 130 trials running window. **C**. The monkey’s choices in a sample sequence of 30 trials. Color intensity indicates signed motion coherence direction, with positive values showing motion toward T1 and negative values motion toward T2. **D - F**. Psychometric curves from one experimental session (the same session as in B-C) computed conditioned on the monkey’s slow choice preference (D), previous choice (E), and previous reward (F). Dots indicate actual data and the solid lines are maximum likelihood fits of logistic functions (see Methods).

### Highly trained monkeys exhibit slow and fast choice fluctuations

Monkeys were extensively trained in the task and showed stable performance prior to neural recordings. Despite their extensive training and trial-to-trial independence of stimulus conditions, both monkeys demonstrated slow fluctuations in their choice preference, where monkeys chose one target more frequently than the other for tens to hundreds of trials before reversing their preference (Figure 1B; black line at the bottom. These fluctuations were spontaneous and could not be explained by fluctuations of motion direction across trials because motion direction was largely balanced in those periods (Figure 1B, top). To further ensure that the slow choice preference fluctuation did not merely reflect random fluctuations arising from spurious unbalance of motion directions and coherence, we repeated our analysis and replicated our results by subsampling trials to equalize the number of trials with stimuli moving toward T1 or T2 for each coherence (Figure 1B and S1A-B blue line; see Methods, Equation 2). Across sessions, the correlation coefficient between slow choice preference fluctuations and signed motion coherence was weak and not statistically significant (mean ± SEM 0.02 ± 0.04, permutation test *p*-value = 0.26). Additionally, the mean auto-correlogram of slow choice preference fluctuations calculated on coherence-balanced trial history showed a statistically significant broad central peak (Figure S1; permutation test, *p*-value < 0.05), indicating the presence of slow fluctuations of response preference, irrespective of fluctuations of stimulus statistics.

On a finer timescale, we also observed that the monkey’s choices were influenced by recent choices and outcomes. As illustrated by the example trial sequence in Figure 1C, the monkey tended to choose the opposite target after error trials in this session. Fluctuations of both slow and fast choice preference were reflected as a shift in the psychometric curve (Methods, Equation 1) when it was calculated conditioned on the direction of slow choice preference (Figure 1D), previous choice (Figure 1E), or recent outcome (Figure 1F).

### Improvement of choice prediction accuracy with slow and fast choice fluctuations

To quantify how monkeys’ decisions were affected by the slow and fast fluctuations of choice preference, we measured whether and how much they would improve the prediction accuracy of upcoming choice beyond that given by motion stimulus alone.

We built a logistic regression model to predict choices based on three variables: stimulus coherence, slow choice preference fluctuation, and fast choice preference fluctuations expressed as a combination of previous choice and reward (Methods, Equations 2-4). The cross-validated model prediction accuracy was assessed using a *leave–one–out* procedure. To measure the role of choice preference fluctuations in determining choices, we compared the prediction accuracy with a baseline obtained by fitting the model using shuffled fast and slow choice preferences across trials. Because shuffling destroys the statistical relationship between current choices and choice preference fluctuations, the comparison with the baseline isolates improvement of the prediction accuracy conferred by the latter. The mean prediction accuracy improvement across all sessions was small but significantly larger than zero (0.016 ± 0.002, one sample *t*-test, *p*-value = 10^−6^; Figure 2A; black bars). Importantly, when we focused only on difficult trials, where the stimulus is less informative and biases could have a larger influence on choice, the improvement of choice prediction accuracy doubled (mean ± SEM: 0.032 ± 0.004; one sample *t*-test, *p*-value = 10^−7^; Figure 2A; open bars). Consistently, the improvement tested only on easy trials was not different from zero (mean ± SEM: 0.0001 ± 0.0002; one sample *t*-test test, *p*-value = 0.69), which is expected because prediction accuracies based on stimulus strength alone are already close to ceiling. Overall, biases had a tangible effect on upcoming choices especially for more difficult decisions.

**Figure 2.**
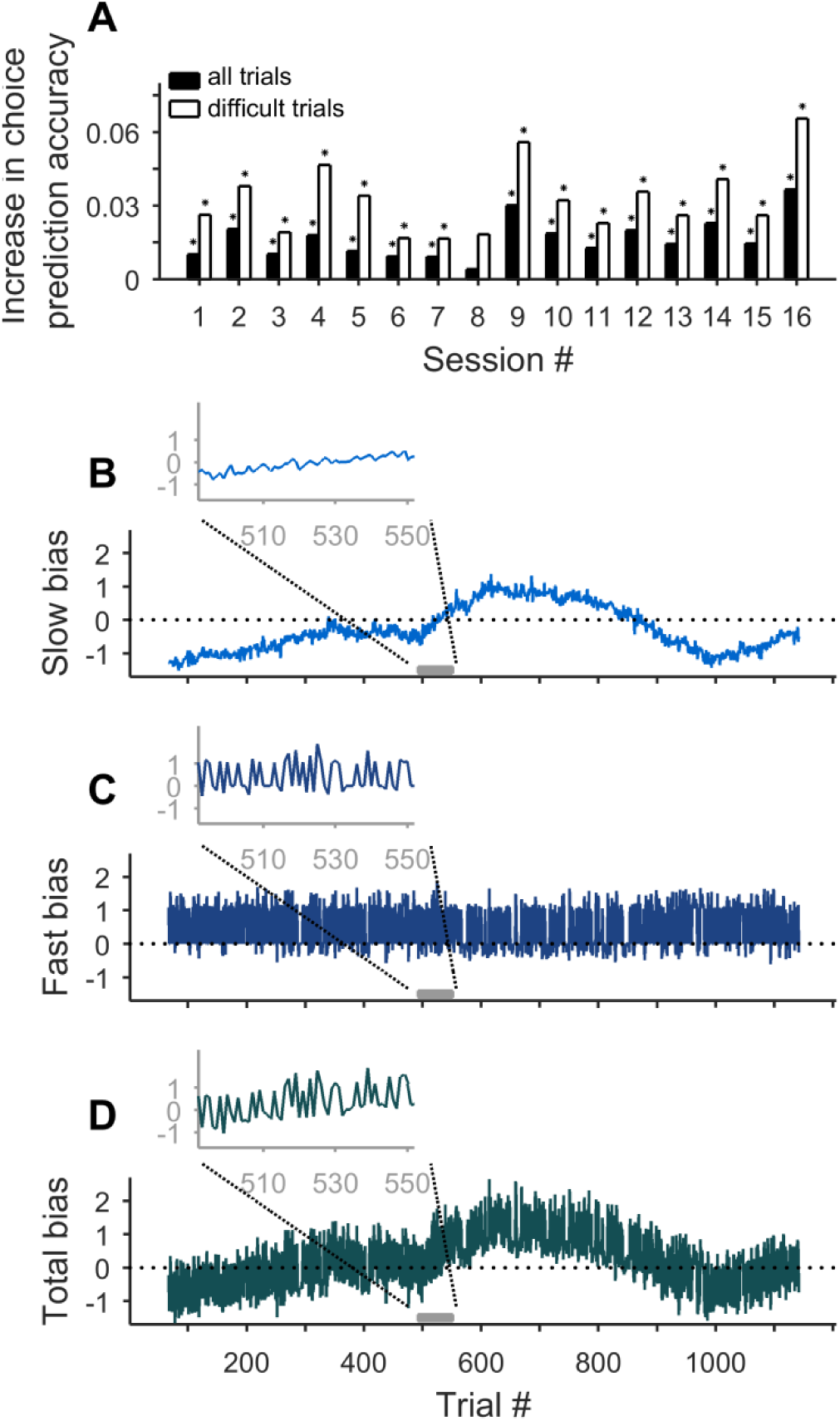
Predicting the monkey’s choices becomes more accurate by using fluctuations of behaviorally-defined choice preference. **A**. Improvement of choice prediction accuracy measured as the difference of the accuracy of a logistic regression model with fast and slow choice preference fluctuations and a reference model in which choice preferences were shuffled across trials. Both models contained stimulus strength (signed motion coherence) as a regressor in addition to the bias terms. Improvement in choice prediction accuracy was higher when computed for difficult trials (white bars, compare to black bars for all trials). Each pair of bars represents a single session. **B - D**. Traces of slow (B, blue), fast (C, dark blue) and total (summed fast and slow, D, green) biases across a sample experimental session. On each trial, biases were computed from the choice preference fluctuations multiplied by their corresponding weights from the logistic regression model. Insets zoom in on a sequence of 60 trials in the session (the grey bar on the x-axis) for better visualization of the dynamics of different types of biases. Note the different time scale between slow and fast biases but their similar contribution to the total bias in terms of their magnitudes.

It is possible to use similar choice prediction models to quantify the temporal extent of fast and slow choice preference fluctuations. For fast choice preference fluctuations, information about choice and reward of the previous trial significantly improved choice prediction accuracy (Equation 5 in Methods, mean difference tested on difficult trials equaled 0.015 ± 0.0006; one sample *t*-test, *p*-value = 3*10^−4^). However, including information about more distant past (from two to five trials back) did no improve the prediction accuracy further (paired *t*-test on prediction accuracies, *p*-value > 0.33; Figure S2A), indicating that in our task, fast choice fluctuations were shaped in a time scale that was not longer than one single trial in the past.

We studied how the size of the trials window used to calculate slow choice preference improved performance of a model that included both slow and fast choice preference fluctuations compared to a model in which only fast choice preference fluctuations was used (in both models coherence was also used as a regressor). If slow fluctuations were defined using trial windows of less than 130 trials, there was not statistically measurable effect (Figure S2B; Equation 4, paired *t*-test on prediction accuracies, *p*-value > 0.063). However, when slow fluctuations were calculated in larger windows (130 - 400 trials), there was a statistically significant increase in prediction accuracy (paired *t*-test rank sum test, *p*-value < 0.05 for 19 cases; Figure S2B). Therefore, we conclude that slow choice preference fluctuations estimated in a window of ∼130 trials and fast choice fluctuations capturing the immediately preceding choice and reward were sufficient to explain the behavioral biases observed in our experiment.

An important question is to know the relative strength of the effect of fast and slow choice preference fluctuations. To compare their effects, we first expressed both types of fluctuations in the same units (log-odd units) by using the logistic regression model described above. Thus, “slow bias” (Figure 2B) was defined as the product of slow choice preference fluctuation and its corresponding weight in the model plus the model offset (see Methods). Similarly, “fast bias” (Figure 2C) was defined as the product of fast choice preference fluctuation and its model weights (see Methods). To compare the relative strength of slow and fast biases on choice prediction, we calculated the mean ratio of the absolute value of each divided by the sums of absolute values of both biases and stimulus strength in log-odds space (see Methods Equation 6). We found that both fast and slow biases were effective in shaping the choice but, on average, the fast bias had a lower impact on choice (mean ± SEM 0.18 ± 0.01) compared to the slow bias (mean ± SEM 0.25 ± 0.01; difference of fast and slow = - 0.07 ± 0.016, one sample *t*-test *p*-value = 0.006).

To capture the overall effect of slow and fast biases on behavior, we defined “total bias” (Figure 2D) on each trial as the sum of slow and fast biases. The total bias corresponds to the single-trial quantification of bias that is central for our analysis. On average, the contribution of total bias to the monkey’s choices was 0.25 ± 0.01 in units of log-odds. For comparison, it was more than twice smaller than the contribution of motion stimulus alone (mean ± SEM 0.58 ± 0.02), reflecting the fact that monkeys made their decision largely based on the presented stimuli but they were also slightly impacted by their current bias.

### Representation of bias in the pre-stimulus neural responses of PAG

Since the slow and fast biases influenced the monkey’s choice (Figure 2A), the information about them should be present in brain regions involved in the decision-making process. An area of interest could be PAG, where neural responses related to the accumulation of evidence have been found (Kiani et al., 2014b; Kim and Shadlen, 1999; Mante et al., 2013). Also, since the observed behavioral biases had a history dependent component, there should be neurons sensitive to the bias even prior to the stimulus presentation. Such a tuning is illustrated in Figure 3A-C, where we show responses of example neurons modulated by slow (Figure 3A), fast (Figure 3B), and total (Figure 3C) biases.

**Figure 3.**
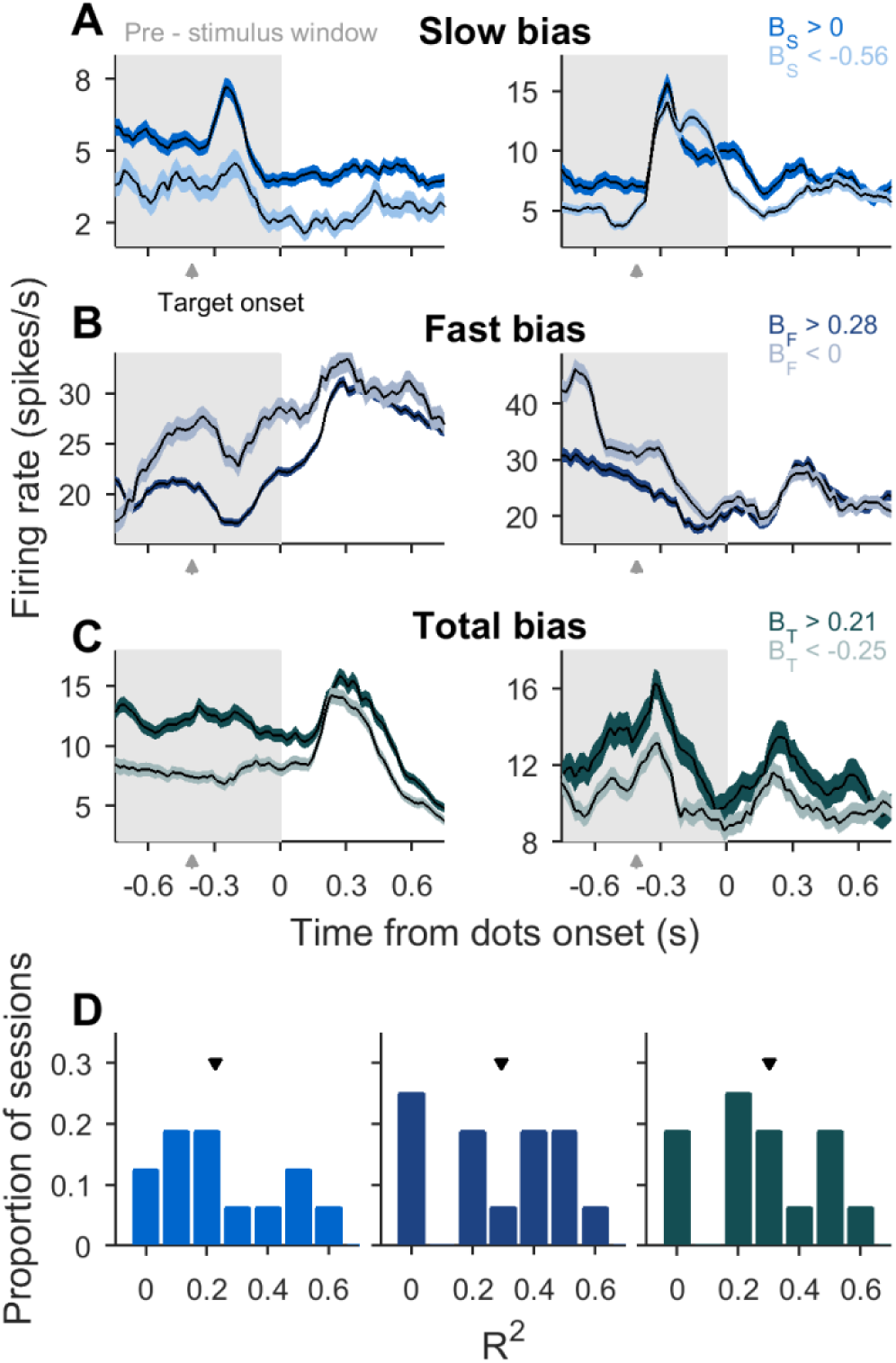
Pre-stimulus responses of prearcuate gyrus neurons carry information about behavioral biases. **A - C**. Peristimulus time histograms (PSTHs) of example cells (left, monkey 1, and right, monkey 2) averaged across trials with high positive (dark) or negative (light) slow (A, blue), fast (B, red), or total (C, green) biases. Specific bias ranges for which trials were averaged (legends) corresponded to the 0.35 (negative) and 0.65 (positive) quantiles of the bias distribution for all sessions pulled together. Shaded areas correspond to SEM. On the x-axis, zero refers to stimulus onset. Target onset time is indicated by an arrow. Spikes were counted in 100 ms moving windows in steps of 20 ms. **D**. Histograms of the coefficients of determination (cross-validated R^2^) of a linear regression model fitted to predict slow (left, blue), fast (middle, red), or total (right, green) biases from the pre-stimulus population activity of prearcuate gyrus cells (T = 800 ms; grey areas in A – C). Arrows indicate mean across sessions (0.23 ± 0.05 for slow; 0.29 ± 0.06 for fast and 0.3 ± 0.05 for total biases respectively; mean ± SEM).

To investigate whether these biases were represented in the responses of PAG neural population, we used a linear model in which each type of bias (slow, fast, or total) was regressed against pre-stimulus activity of simultaneously recorded PAG neurons (Equation 8; spike counts calculated in 800 ms window prior to stimulus onset). The analyses were performed on the top variance-predictive PCA components of the neural population responses that collectively explained 50% of the variance (Equation 7). Here, we used PCA to reduce overfitting (Mante et al., 2013; Yu et al., 2009), but qualitatively similar results were obtained without dimensionality reduction. All three biases were significantly represented in the pre-stimulus responses of PAG population (Figure 3D; mean of cross–validated *R*^*2*^ across sessions was equal to 0.23 ± 0.05, 0.29 ± 0.06 and 0.30 ± 0.05 for *B*_*s*_, *B*_*f*_ and *B*_*t*_ respectively, permutation test, *p*-value = 0.001 in all three cases).

Consistent with a representation of the fast bias, PAG activity prior to the stimulus onset represented previous choice and reward, which, together, defined the fast bias. Examples of four neurons for which firing rates were modulated by previous choice or reward are shown in Figure S3 (A and C respectively). In all cases, the response modulation toward previous choice or recent outcome persisted through the pre-stimulus period.

Fitting a logistic regression model to the firing rate of the population of simultaneously recorded PAG neurons (Equation 9 for *n* < 0) revealed that previous choices could be decoded up to three trials back in the past (Figure S3B; means ± SEMs: 0.52 ± 0.005; *p*-value = 0.006 for *n* = −3; 0.53 ± 0.006; *p*-value = 0.0007 for *n* = - 2 and 0.81 ± 0.02; *p*-value < 10^−10^ for *n* = −1, one sided *t*-test). A similar model (Equation 9 for *n* < 0) could predict the outcome (reward) of the preceding trial (Figure S3D; mean ± SEM 0.81 ± 0.01; paired *t*-test, *p*-value 0.0002), but not further back.

### Predicting choices from pre-stimulus neural responses of PAG

Given that slow and fast biases influenced monkeys’ decisions (see Figure 2A) and that they were represented in the pre-stimulus PAG activity (see Figure 3), we asked whether pre-stimulus activity was also predictive of monkeys’ upcoming choices. Figure 4A shows two example units. One of the units (Figure 4A, left) had a higher firing rate for choosing the T2 target and the other unit had a higher firing rate for the opposite target (Figure 4A, right). Importantly, both units represented the upcoming choice even prior to the stimulus presentation (grey areas), matching the representation of bias in the neural population.

**Figure 4.**
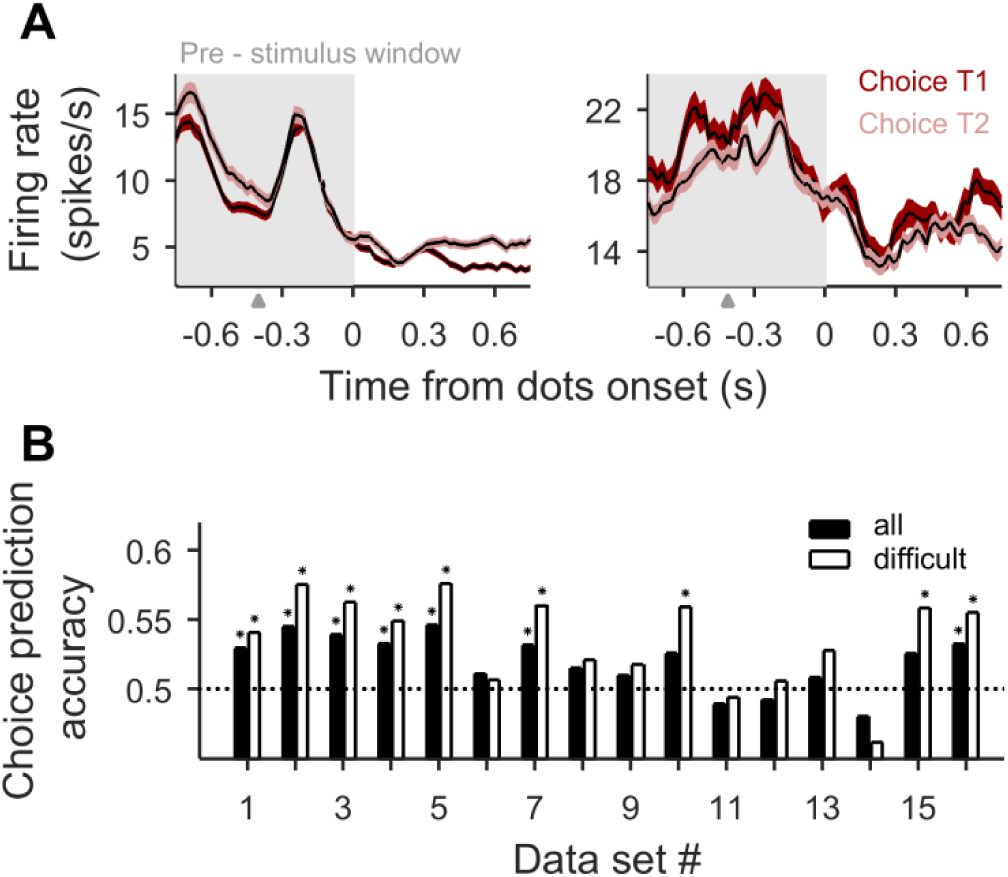
Pre-stimulus activity of prearcuate gyrus neurons predicts upcoming choice. **A**. Mean firing rate of two cells (from two monkeys) averaged across trials with T1 (dark) or T2 (light) choices. Shaded area corresponds to SEM. Arrow on the x-axis indicates target onset. Firing rates were calculated in a 100 ms windows moved in steps of 20 ms. **B**. Cross-validated prediction accuracy of a logistic regression model for predicting upcoming choice from the pre-stimulus activity of simultaneously recorded neurons in prearcuate gyrus (window size, 800 ms; grey area in A, PCA dimensionality reduction). Accuracy was higher when assessed for difficult trials only (white bars). Choice prediction accuracy was above chance level (0.5, dotted line) for seven or nine sessions when calculated for all or difficult trials respectively (*, permutation test; *p*- value < 0.05). Mean across session was equal (0.52 ± 0.005, *p*-value = 0.001 for all trials; 0.54 ± 0.008 *p*-value = 0.001 for difficult trials mean ± SEM *p*-values from permutation test).

To quantify this activity modulation at the population level, we fit a logistic regression model to predict upcoming choices using the PCA-dimensionality-reduced PAG population responses in the 800 ms before stimulus onset (Equation 9 for *n* = 0). For many sessions (44%; 7 out of 16), the cross–validated prediction accuracy was above chance level (0.55) (Figure 4B, black bars; mean across sessions, 0.52 ± 0.005, permutation test *p*-value = 0.001). Similar to the behavioral model, the cross-validated prediction accuracy was higher when we focused on difficult trials (white bars; 0.54 ± 0.008 permutation test, *p*-value = 0.001), and not significantly different from chance level for easy trials (mean ± SEM across sessions 0.5 ± 0.004 permutation test, *p*-value = 0.31).

### Prediction of choices based on pre-stimulus neural activity is due to the representation of slow and fast biases

A key question is whether choice predictive neural responses prior to stimulus onset are due to the representation of the fast and slow biases that we have defined behaviorally. One possibility is that encoding of our behaviorally-defined biases fully explains the representation of choice prior to the stimulus onset, which would imply that total bias and choice representations are “aligned” in neuronal activity space. Alternatively, choice predictive neural responses could arise from factors not fully captured by fast and slow biases, which would cause misalignments between choice and total bias representations (see Methods, Equation 13). To differentiate these two possibilities, we asked if the neural representation of biases was as predictive of behavior as the neural representation of choice.

For each trial, we used the remaining trials in the session to find the best hyperplanes that explained the choice (Equation 9) and total bias (Equation 8) based on pre-stimulus responses (Figure 5A). Then, we calculated the distance of pre-stimulus responses of the left-out trial from those two hyperplanes (*d*_*choice*_and *d*_*bias*_). If the pre-stimulus choice and bias representations were aligned, predicting the upcoming choice based on *d*_*bias*_ would be as accurate as using both *d*_*choice*_and *d*_*bias*_ (Equations 10 and 11). Indeed, this was what we observed. Across sessions, the difference in predicted accuracy was negligible (Figure 5B; mean ± SEM, 0.002 ± 0.003) and not significant (paired *t*-test, *p*-value = 0.42), suggesting that the neural representation of total bias was functionally indistinguishable from the neural representation of choice prior to stimulus onset. Consistent with these results, predicting the choice based on the sign of *d*_*bias*_ or *d*_*choice*_was comparable, with slightly better accuracies for *d*_*bias*_ (Figure. 5C; mean prediction accuracy based on choice hyperplane, 0.52 ± 0.005; permutation test *p*-value = 0.001); based on bias hyperplane 0.54 ± 0.004; permutation test *p*-value = 0.001), further supporting our conclusion.

**Figure 5.**
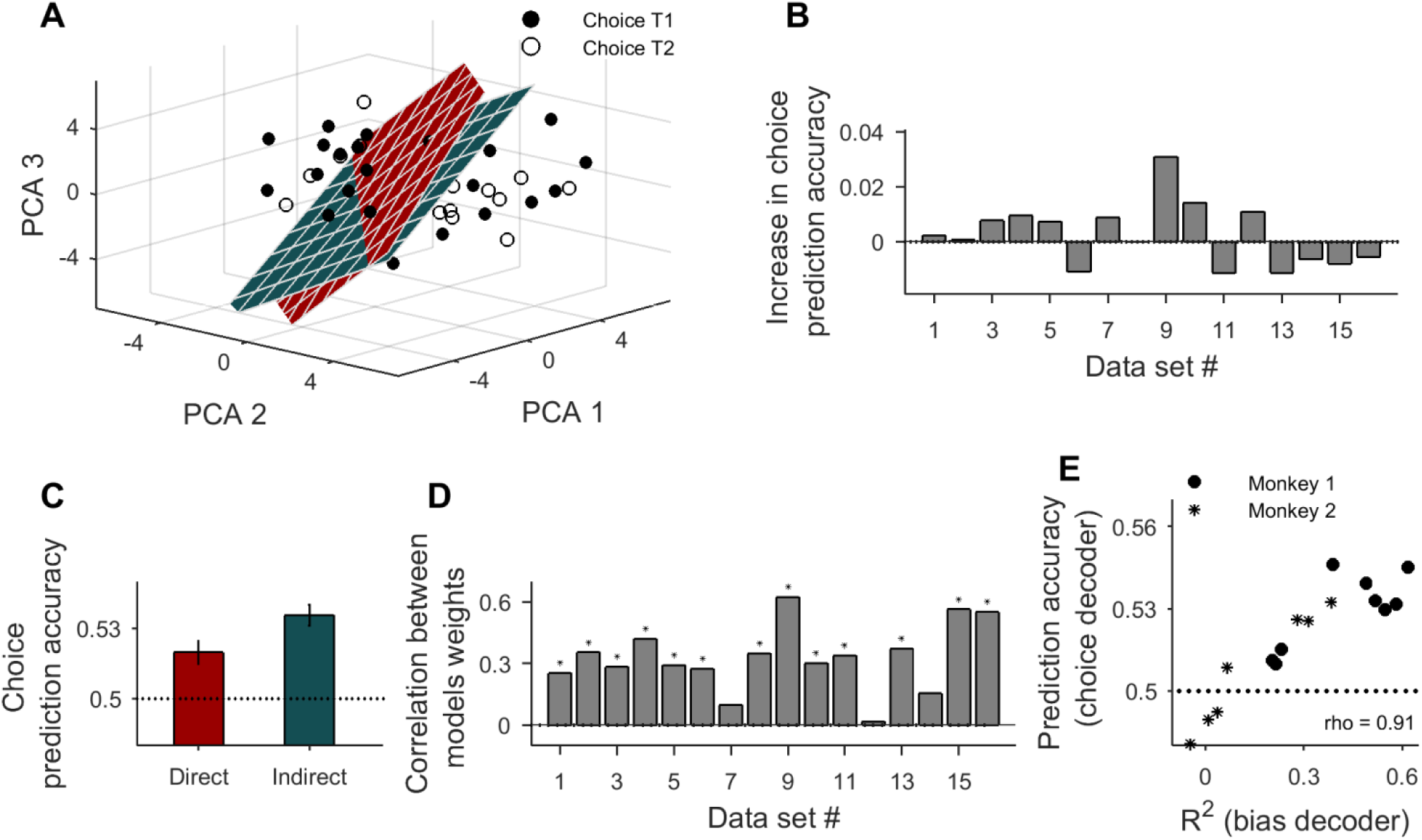
Choice predictive power of pre-stimulus PAG neural responses is related to the representation of total bias. **A**. The total bias (green) and choice (red) decoders, corresponding to two discriminant hyperplanes that split neuronal state space in two regions for T1/T2 choices or positive and negative total bias, were trained using the same pre-stimulus activity of PAG neuronal populations. The panels show example data points (40 trials) where each dot represents the pre-stimulus population firing rates of a trial projected on the first three PCA dimensions. Filled circles correspond to T1 choices and hollow circles to T2 choices. Although the bias decoder has been trained to predict biases, it can be used “indirectly” to predict choices because the sign of the bias indicates tendency toward a choice in each trial (“indirect method”). In contrast, the “direct method” uses the choice decoder to predict choices. **B**. Difference in choice prediction accuracy between a model including the distances from neuronal activity to the choice and bias hyperplanes as predictors, and a model with the distance to the bias hyperplane as the only predictor. The mean prediction accuracy difference between the two models was equal 0.002 ± 0.003 and not significant (paired *t*-test, *p*-value = 0.42) **C**. Experimental data. Choice prediction accuracies from the direct and indirect methods were comparable, suggesting that the bias and choice decoders are aligned as illustrated in (A). As in (B), choice decoding was based on PAG activity from a 800 ms window before stimulus onset (0.5 chance level marked by dotted line; mean ± SEM calculated across sessions, PCA dimensionality reduction) **D**. Consistent with an alignment of the bias and choice hyperplanes, the correlation coefficient between weights of choice and bias decoder hyperplanes (y-axis) was significantly positive for most of the sessions (* permutation test, *p* - value < 0.05). **E**. Across sessions choice prediction accuracy (direct method) correlated with the strength of the total bias representation (defined as the R^2^ of the total bias linear regression model; *corr* = 0.91, *p* - value = 0.0004). Each point represents a single session (dots or stars for Monkey 1 or 2, respectively).

Additional insight about choice predictive neural responses prior to stimulus onset is gained from comparing the geometry of choice and bias decoder hyperplanes. Vectors in high-dimensional spaces tend to be orthogonal (Hall et al., 2005). However, if the representation of choice and bias are functionally aligned, one would expect that the angle between the norms of their respective hyperplanes is less than 90 deg. In fact, we found that the weight vectors that defined the norm of the choice and bias hyperplanes (*β*_*j*_ in Equation 8 and *α*_*j*_ in Equation 9) were positively correlated and the correlation coefficients were significant for the majority of sessions (Figure 5D; n = 13, permutation test *p*-value < 0.05; across session mean ± SEM, 0.33 ± 0.04; permutation test *p*-value = 0.001), supporting the notion that the hyperplane norms were not orthogonal.

Further supporting the alignment hypothesis, we found that sessions with stronger representation of the total bias (higher cross-validated R^2^) in pre-stimulus activity of PAG also had a higher cross-validated choice prediction accuracy (Figure 5E; Pearson correlation coefficient 0.91 *p*-value = 10^−7^). These results suggest that across-session variability in choice predictive power could be explained by the across-session variability in the representation of total bias.

### The integration of bias into the accumulation of evidence

Given the presence of the bias signal prior to stimulus presentation, the question arises of whether and how this bias impacts the accumulation of the sensory evidence during the stimulus-viewing period. Our result about the alignment of bias and choice decoders before stimulus onset suggests that the bias could be implemented as an initial offset in baseline activity before accumulation of sensory information begins in each trial. This initial offset is best visible in the neuronal activity axes where the decision is encoded (orthogonal to the choice decoder hyperplane shown in Figure 5A). Thus, we plotted how the decision variable (projection of neuronal activity onto the choice axes) evolved over time (Figure 6, see Methods Equation 14).

**Figure 6.**
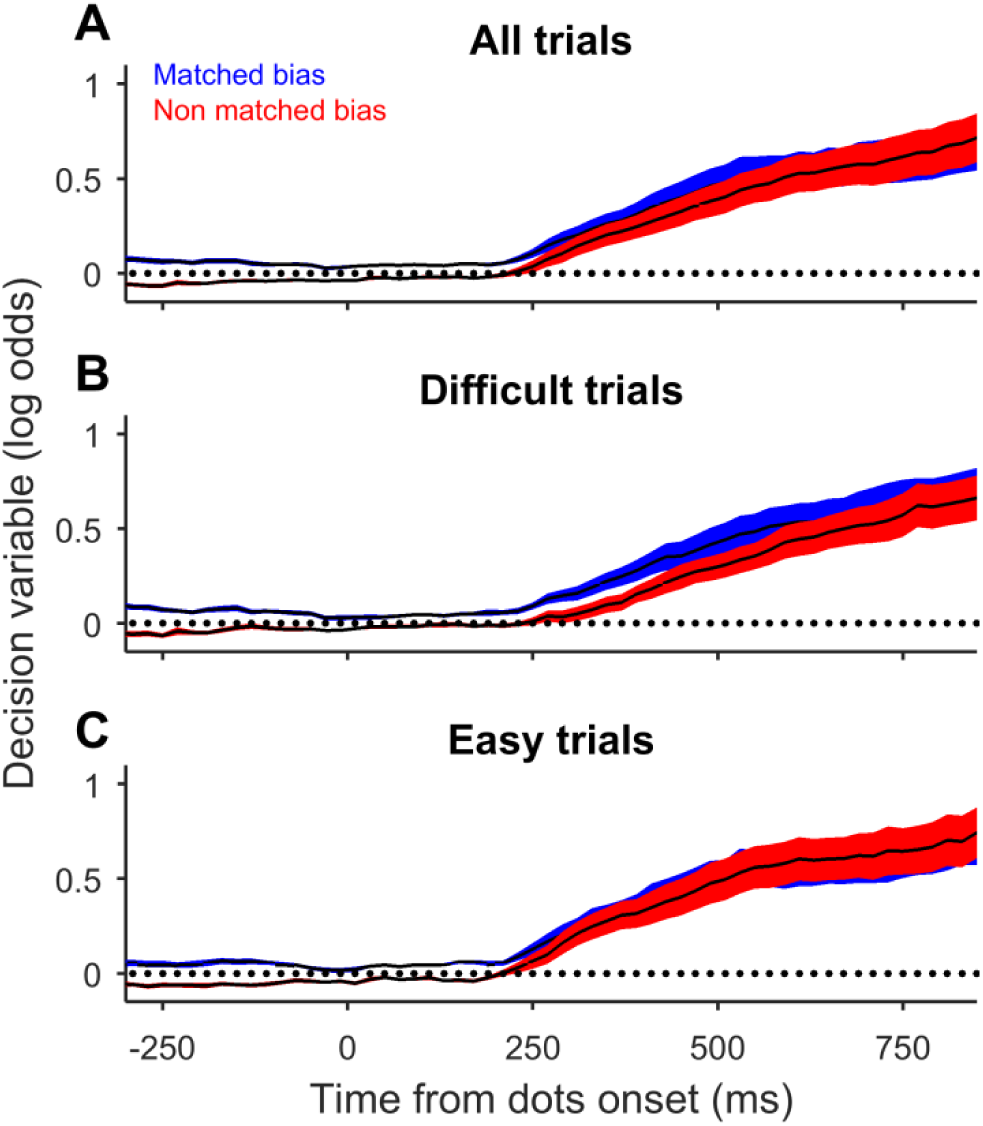
Total bias acts as an initial offset of the decision variable before accumulation of motion information begins. **A - C**. Instantaneous decision variable (DV) from the logistic regression model of choice. The DV is averaged across trials in which the total bias was aligned with the monkey’s choice (blue) or trials in which the total bias was against the final choice (red, non-matched). Different panels show the average DVs for all (A), difficult (B) or easy (C) trials. The DVs were calculated based on 100ms of dimensionality-reduced neural responses centered at each time (PCA axes for dimensionality reduction were the same as those used in earlier figures and calculated for the 800ms window before stimulus onset). The analyses window was moved in steps of 20 ms. Shading indicates SEM.

We found, consistent with our expectations, that there was an offset in the decision variable before stimulus onset. The offset was positive when the bias favored the final choice (Figure. 6A, blue line) and negative when the bias was against the final choice (red). This offset was roughly constant for the whole duration before stimulus onset and persisted during the first few hundreds of milliseconds of stimulus viewing period, when the decision was formed. Interestingly, the offset lasted longer for more difficult stimuli (Figure 6B), where monkeys integrated the sensory evidence longer. However, toward the end of the stimulus presentation, the offset vanished because the monkey had likely reached a decision on the majority of trials. Our results suggest that the initial bias was integrated into the decision-making process and contributed to the formation of the choice.

The initial offset and final convergence of the decision variables for positive (Figure 6A, blue line) and negative (red line) biases are consistent with predictions of bounded evidence-accumulation models for the decision-making process (Link, 1992; Ratcliff and Smith, 2004; Shadlen and Kiani, 2013). The gradual dynamics of the decision-variables and larger and longer-lasting effects of offset for weaker motion stimuli (compare Figure 6B and 6C) is compatible with these models too. This is because the accumulation of sensory evidences is slower for weaker stimuli, and the decision variable takes longer to hit the decision bound. This slower ramping lets the bias-induced offset (blue and red) survive longer during stimulus viewing. In contrast, sensory evidence accumulates quickly on easy trials, causing accelerated convergence of the decision variables and leaving minimal room for the bias-induced offset to influence the final decision.

Another interesting feature of the neurally inferred decision variable is that it continues to rise after the convergence of positive and negative bias curves in Figure 6. Because our model is designed to predict the choice (Equation. 14), the magnitude of the model decision variable is influenced by any factor that improves its accuracy. Those include neural responses that may represent only the final choice but not necessarily the decision-making process that leads to the choice. Such choice-related responses have been shown to emerge in motor-planning regions toward the end of the stimulus viewing period in the dots task (Peixoto et al., 2018), and could be responsible for additional rise of the model decision variable after 700ms from stimulus onset, when the initial bias is no longer represented.

## Discussion

We have studied the dynamics and neuronal representation of biases in highly-trained monkeys performing a direction discrimination task while recording simultaneous responses of hundreds of neurons in the prefrontal cortex. Despite trial-by-trial independence of the stimulus direction, monkeys exhibited weak but measurable suboptimal behavioral biases. Observed biases emerged at two distinct time scales. The slow bias, reflecting the monkey’s preference towards one of the targets, fluctuated at a time scale spanning tens to hundreds of trials. The fast bias was shaped by the choice and outcome of only the preceding trial. Together these biases improved prediction accuracy of upcoming choices beyond that given by stimulus alone. As expected, this increase was higher on trials with weak stimuli. Further, we found that pre-stimulus population activity of prearcuate gyrus represented the fast and slow biases. The same activity was also predictive of the monkey’s upcoming choices. Critically, the axes that represented bias and choice in the neural population state space were similar; suggesting that choice-prediction power of pre-stimulus prefrontal activity was largely due to the representation of the fast and slow biases. Conditioned on behavioral biases, we demonstrated that biases were incorporated into the decision-making process as an offset of baseline activity that persisted throughout the integration of sensory evidence during motion stimulus presentation.

To make optimal decisions one should take into account all available and relevant information. This idea is expressed in Bayesian decision theory where choices depends on both, the current sensory information and prior expectations (for review see (Summerfield and de Lange, 2014)). Such prior expectations might reflect for example the previously learned statistics of the stable environment and as such ease correct decisions especially when sensory evidence is ambiguous (Hanks et al., 2011; Rao et al., 2012). In contrast, when environment is unpredictable, prior expectations do not provide additional useful information into decision at hand but instead might lead to suboptimal behavioral biases that potentially reduce accuracy. Our task was designed to make prior history uninformative, and thus all biases that we observed despite extensive monkey’s training could not arise from a reward optimization strategies. Rather, they provide an unique opportunity to explore innate mechanisms that shape history-dependent biases.

In perceptual and value based decision making tasks one particular type of biases refers to the fact that current choices can depend on previous history of stimuli, rewards and choices even in the conditions where such dependence is not relevant for the task – so-called sequential biases (Abrahamyan et al., 2016; Akrami et al., 2018; Busse et al., 2011; Eskandar and Assad, 1999; Fritsche et al., 2017; Hermoso-Mendizabal et al., 2020; Lueckmann et al., 2018; Nogueira et al., 2017; Padoa-Schioppa, 2013; Shadlen and Newsome, 2001). Such history dependences can span from one trial up to several trials in the past (Akrami et al., 2018; Busse et al., 2011; Gold et al., 2008b; Hermoso-Mendizabal et al., 2020; Lueckmann et al., 2018; Nogueira et al., 2017). However, only a couple of studies have reported the presence of separate time scales in sequential biases. One such example refers to the study where monkeys performed perceptual decision task and during training exhibited intrinsic slow (tens of trials) and fast (previous few trials) choice sequential biases (Gold et al., 2008b). As the slow bias component decreased with training, it is likely that it was a byproduct of learning. Fast and slow bias components have been shown to depend also on sensory components of the previous history (Akrami et al., 2018). Our results add to that body of research and constitute a rather unique example of such biases in highly trained primates performing unstructured tasks in which past trails are irrelevant for choice at hand. As it has been suggested, the existence of such biases might be a byproduct of a priori adaptive mechanisms that take advantage of stability of natural environment to make faster and more accurate decisions (Summerfield and de Lange, 2014). However, in our task biases, while having an impact on choice, they have very little impact on task performance.

Given that history-dependent biases impact choices, they should be represented in the form of choice-predictive neuronal activity in brain regions that represent both the bias and the decision-making process. Such choice predictability has been described in the pre-stimulus activity of LIP neurons in the seminal work of Shadlen and Newsome (Shadlen and Newsome, 2001). However, as the authors did not investigate history-dependent biases, it was unclear what the source of this choice predictability was. Several studies investigated whether choice predictability reflects past history of stimuli, choices and outcomes (Akaishi et al., 2014; Akrami et al., 2018; Bonaiuto et al., 2016; Eskandar and Assad, 1999; Fischer and Whitney, 2014; Lueckmann et al., 2018; Nogueira et al., 2017; Shadlen and Newsome, 2001; Williams et al., 2003; Wyart and Tallon-Baudry, 2009), Although recently it has been demonstrated that choice-predictive signals in visual cortex could be partially accounted for by previous history (Lueckmann et al., 2018), the remaining unexplained residuals probably reflected unmeasured biases at the behavioral level that are nevertheless measurable at the neuronal level. At the behavioral level, our results go beyond these results by providing a single-trial quantification of bias that seems to exhaust all biases that are linearly decodable from neuronal population activity in PAG. That allowed us to demonstrate that choice predictive power of prefrontal cortex pre-stimulus activity can be explained by history dependent biases. While it is still possible that other biases exist in the monkeys’ behavior, their temporality or non-linearity make them hard to detect and measure.

Several models have proposed how biases might mechanistically combine with sensory information to form the final decision (Bogacz et al., 2006; Drugowitsch and Pouget, 2012; Drugowitsch et al., 2019; Gold and Shadlen, 2007; Rustichini and Padoa-Schioppa, 2015). One of the predictions is that biases can act either as an offset or as a change in the slope of sensory information accumulation (Drugowitsch et al., 2019; Gold and Shadlen, 2007; Shadlen and Kiani, 2013; Urai et al., 2019) (but see (Sohn et al., 2019)). Previous experimental work provided evidence in favor of the offset hypothesis (Gold et al., 2008b; Shadlen and Newsome, 2001). For instance, conditioning neuronal responses on the final choice Shadlen and Newsome demonstrated an offset in pre-stimulus LIP neuronal activity that persisted during the stimulus presentation period (Shadlen and Newsome, 2001). However, since the choice is a combination of bias and stimulus evidence integration, whether similar offsets in accumulation of evidence would be observed when directly conditioning on bias has remained unsolved. Here we provide this missing evidence and demonstrate an offset in pre-stimulus neuronal activity between trials in which behavioral bias matched or mismatched final decision.

To conclude, we have provided neuronal evidence that behavioral biases and choices are represented in the same neuronal circuits along similar directions of activity state space. The implications of these results can be multifarious. For instance, in a speculative vein, the fact that biases are directly incorporated into the decision process as an offset, just as veridical information would do, could speak about why it is so difficult to eliminate deleterious biases from our daily life behavior (Gigerenzer, 2008; Kahneman, 2011), and it is in line with current work on decision making proposing bottlenecks in sensory (Moreno-Bote et al., 2014) and value-based processing (Hayden and Moreno-Bote, 2018).

## Acknowledgements

We would like to thank to Mohamad Saleh Esteki, William Newsome, Ramon Nogueira, Gouki Okazawa and Jacob Yates for fruitful discussions about the project. Also we would like to acknowledge the funding agencies: GM is supported by IJCI-2014-21937 from MINECO (Spain); RK is supported by the Simons Collaboration on the Global Brain (542997), McKnight Scholar Award, Pew Scholarship in the Biomedical Sciences, and National Institutes of Mental Health (R01 MH109180-01); RM-B is supported by BFU2017-85936-P from MINECO (Spain), the Howard Hughes Medical Institute (HHMI; ref 55008742), an ICREA Academia award, and the Bial Foundation (grant number 117/18).

## Author contributions

GM, RK and RMB designed the project and analyses; RK provided the data (adopted from (Kiani et al., 2015)); GM analyzed the data; GM, RK and RMB wrote the manuscript.

## Methods

### Experimental Procedures

We recorded extracellular activity from populations of neurons in the prearcuate gyrus (PAG) of two macaque monkeys performing a direction discrimination task. All training, surgery, and recording procedures conformed to the National Institutes of Health Guide for the Care and Use of Laboratory Animals and were approved by Stanford University Animal Care and Use Committee.

### Behavioral Tasks

The direction discrimination task is illustrated in Figure 1A. Each trial began when a fixation point (FP; 0.3° diameter) appeared at the center of the monitor. The monkey was required to fixate within ±1.5° of the FP. Afterwards two targets (T1 and T2) appeared. In 11 sessions, the targets were placed on opposite sides of the screen. In the remaining five sessions, both targets were placed contralateral to the recorded hemisphere. After a short delay (400 ms for 15 data sets and 500–1500 ms for one data set, median 876 ms), a patch of randomly moving dots was shown at the center of the screen for 800 ms. The fraction of coherently moving dots (stimulus strength or coherence) defined the difficulty of a given trial (Britten et al., 1992; Kiani et al., 2008; Shadlen and Newsome, 2001). The motion direction and strength were chosen randomly on each trial from a set of predefined values. The coherent motion direction could be toward one target or the other. Coherence ranged from 0 to 0.8. We use signed motion coherence, C, to specify motion direction and coherence using one number, where positive values indicate motion toward T1 and negative values motion toward T2. The coherence range was tailored for each monkey to obtain the full range of performance accuracy from the chance level to nearly perfect. The stimulus was followed by a delay period of variable duration (302–1478 ms, median = 758 ms) randomly selected on each trial. At the end of the delay period, FP disappeared (Go cue) and the monkey had to report the perceived direction of motion by making a saccade towards the corresponding target and maintaining gaze on the target until the trial outcome was revealed. Correct choices were rewarded with a drop of juice. For the zero coherence trials, choices were rewarded randomly with a probability of 0.5. Throughout the session the eye position was monitored at 1 kHz with a scleral search coil (CNC Engineering, Seattle).

### Neural Recording

We recorded extracellular activity of a population of PAG neurons using 96 channel microelectrode arrays (Blackrock Microsystems, Salt Lake City; electrode length = 1.5 mm; spacing = 0.4 mm; impedance ∼ 0.5 MΩ), while monkeys performed the behavioral task. The electrode array was implanted anterior to the concavity of the arcuate sulcus and posterior to the tip of principal sulcus. Neural signals were saved online with 30 kHz sampling rate and spike waveforms were sorted offline (Plexon Inc., Dallas). Recording artifacts simultaneously occurring in a large number of channels were removed using customized algorithms. We identified 169–250 single and multi-units in each session (median = 220). We use the term ‘‘units’’ to refer to both well-isolated single neurons and multi-units. The data sets analyzed in the present study included 9 and 7 recording sessions from monkeys 1 and 2, respectively. These sessions were chosen based on three factors: large number of trials per session (>1000), high quality of recordings, and large number of units. Although the position of the electrode array could not be changed by the experimenter after implantation, the recorded units could change from one session to another, presumably due to small movements of cortex relative to the array. The analyzed data were published previously (Kiani et al., 2014b, 2015) in the context of different scientific questions.

Due to the length of the experimental sessions and the large size of the datasets, they were saved as multiple small files, each containing data from an experimental block of more than a hundred trials. These data files were concatenated offline at the end of the session. For 11 of the recording sessions, online spike detection thresholds varied in different blocks of the experiment, causing non-stationary baseline firing rates across the session. To make certain that our analyses were not affected by this non-stationarity, we z-scored firing rates in each block after removing the first and last three trials in the block (median number of removed trials per session, 18, range, 6-30). The z-scored firing rates were concatenated across the session and used in the session by session analyses explained below. Similar but noisier results were obtained when the blocks were analyzed separately without z-scoring.

### Behavioral Data Analysis

Psychometric curves were defined as the fraction of trials in which the monkey chose target T1 for a given signed motion coherence, *C*, for each session. We fit a logistic function to the psychometric curve:

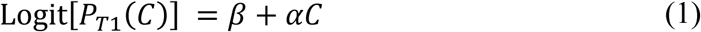

where *P*_*T*1_ is the probability of choosing target T1, and *α* and *β* are model parameters, describing sensitivity and overall bias, respectively. We used a maximum-likelihood fitting procedure for Equation 1 and all subsequent models in the paper.

We use the psychometric function of each session to define difficult and easy motion strengths. Difficult motion strengths were those associated with lower than 75% accuracy. Easy motion strengths were those associated with accuracy equal or higher than 75%.

### Modeling behavior

We hypothesized (and confirmed below) that behavior is influenced by past history of choices at two different time scales: long timescale changes in target preference varying over tens to hundreds of trials, and short timescale preference shaped by action and reward history in a few previous trials.

We defined the slow timescale fluctuations of choice preference based on the frequency of choosing targets T1 and T2:

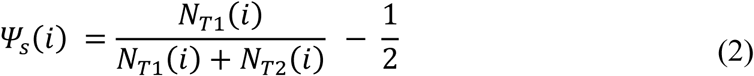

were *N*_*T*1_(*i*) and *N*_*T*2_(*i*) correspond to the number of T1 or T2 choices, respectively. *N*_*T*1_(*i*) and *N*_*T*2_(*i*) were computed for a group of trials around *i* : [*i* − *W*/2, *i* − 2] ∪ [*i* + 1, *i* + *W*/2]. Trials *i* and *i*-1 were excluded from calculation of *ψ*_*s*_(*i*) to avoid confounds and any overlap with the variables that will be used to define the fast timescale fluctuations of choice preference (see below). We tested various trial windows, *W*, as explained below. To ensure that *ψ*_*s*_(*i*) reflected spontaneously generated choice preference and not random fluctuations in the stimulus history, we subsampled trials in the analysis window to balance the number of trials for each signed coherence. Excluding trials after the *i*th trial — making the definition of *ψ*_*s*_(*i*) causal — did not qualitatively change our results.

We defined fast time-scale changes of choice preference based on immediately preceding trials. Past studies suggest that the outcome of decisions before trial *i* can influence the choice on current trials *i* (Abrahamyan et al., 2016; Busse et al., 2011; Hermoso-Mendizabal et al., 2020). We use three indicator variables to define different combinations of choice and outcome for each preceding trial. For trial *i-n*, where *i* indicates the current trial and *n* indicates how many trials back in the past are considered, the vector of indicator variables are:

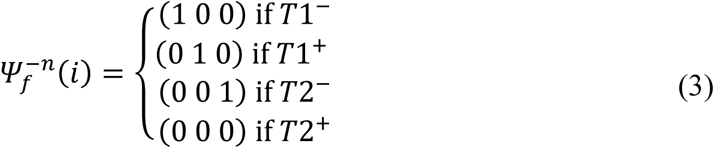

where T1 and T2 indicate the two choices, and +/- indicate the two possible outcomes (rewarded/unrewarded) of trial *i - n*. Here and further in the text we use bold symbol notation to denote vectors.

To test the effect of fast and slow timescale choice preference on the monkey’s behavior, we used a logistic regression model (Hastie et al., 2001) that included both the motion strength and direction from the current *i*^*th*^ trial (signed coherence *C*), together with the slow and fast choice preference fluctuations:

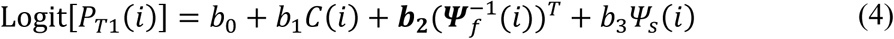

where *P*_*T1*_(*i*) is the probability of a T1 choice in trial *i*. The predictor 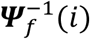 defines the fast choice preference fluctuations, which are shaped by choice and reward on the previous trial (Equation 3), and *ψ*_*s*_(*i*) is the slow choice preference fluctuation that varies at time scales much larger than the immediately experienced history (Equation 2). Here the model weights *b*_0_, *b*_1_ and *b*_3_ are constants and ***b*_2_** is a row vector composed of three constants.

The model in Equation 4 was cross-validated using a leave-one-out procedure: the model was fit to all trials except for a held-out trial and its preceding trial, which was used for estimating the fast choice preference for the held-out trial. We used the model parameters to predict the probability of a T1 choice in the held-out trial. The procedure was repeated for all trials in the experiment. For each training set we balanced the number of T1 and T2 choices by randomly removing trials corresponding to the surplus choice. We considered that the model prediction was correct if it gave a higher probability to the target chosen by the monkey. The overall prediction accuracy of the model was calculated for all trials in a session, or sub-groups of easy and difficult trials.

We used the model described in Equation 4 to estimate fast and slow biases on each trial of each session. The “*fast bias”* at trial *i* was calculated as 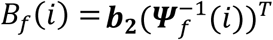. The “*slow bias”* was calculated as *B*_*s*_(*i*) *= b*_0_ + *b*_3_*ψ*_*s*_(*i*). We defined the “*total bias”* as the sum of the fast and slow biases, *B*_*t*_(*i*) *= B*_*f*_(*i*) + *B*_*s*_(*i*). The advantage of defining fast and slow bias effects on the current choice using the parameters obtained by the logistic regression model is that all variables are measured using log-odds, thereby allowing a direct comparison between the strength of the three variables on choices.

The significance of the effects of slow and fast choice preference fluctuations on choices was assessed by shuffling 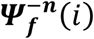 and *ψ*_*s*_(*i*) independently across trials and fitting Equation 4 to this shuffled data. We repeated this process 1000 times to calculate the distribution of prediction accuracy for the shuffled data, corresponding to the null hypothesis distribution. As in our shuffling procedure we kept the relation between the monkey’s choice and motion coherence intact, the null hypothesis distribution was centered on the baseline prediction accuracy given solely by the stimulus direction and strength. We subtracted that mean from the model prediction accuracy to calculate the accuracy improvement conferred by adding the monkey’s fast and slow biases into the logistic regression model. The *p*-value for the significance of this improvement was calculated as the fraction of shuffles for which the prediction accuracy was higher than or equal to the prediction accuracy for the unshuffled data (one-tailed permutation test). To test whether the mean improvement across all the behavioral sessions was different from zero, we used a paired *t*-test between the predicted and mean shuffled accuracies across sessions.

To select the time scale of the slow fluctuations of choice preference, *ψ*_*s*_(*i*), we tested a wide range of *W* from 20 to 500 trials in steps of 10 trials. A wide range of window sizes provided choice prediction accuracies significantly higher than a model without slow choice preference. We chose the shortest window in a consecutive set of significant window sizes (*W* = 130; Figure S2). This window was used for calculation of slow choice preference in all subsequent analyses, but the results were qualitatively similar for other significant window sizes in Figure S2. The time scale of fast bias was chosen using a similar procedure by progressively including 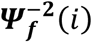 to 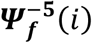 in the model:

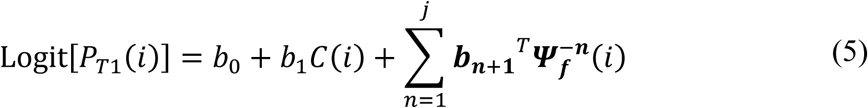

where j = {1, …, 5}. Because choice prediction accuracy did not show tangible improvement by these extensions, we limited our definition of fast bias to the immediately preceding trial, as in Equation 4.

To compare the strength of each bias on the decision, we used the model in Equation 4 to define the motion coherence in log-odds space of choice as *M*_*c*_(*i*) *= b*_1_*C*(*i*). Next for each session separately, we defined the total choice predictive power in *i*^th^ trial as *E*_*t*_(*i*) *=* |*M*_*c*_(*i*)| + |*B*_*f*_(*i*)| + |*B*_*s*_(*i*)|. The impact of each bias (*I*_*x*_) on the decision, (where *x* ∈ {*s, f, t*} stands for slow, fast or total bias) was defined as:

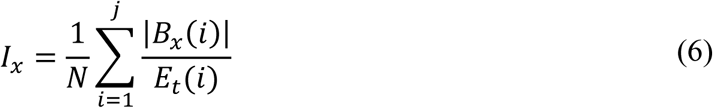

To evaluate whether variations of total bias, *B*_*t*_, correlates with the monkey’s accuracy, we measured average accuracy in windows of 130 trials. Accuracy was based on a subsampled group of trials to balance motion coherences. We used 130 trials to match the time window in which total bias was calculated. Next we calculated Pearson correlation between absolute value of total bias and accuracy. To assess the significance of Pearson correlations, we used a permutation test by randomly shuffling rewarded trials and recomputing accuracy and its correlation with absolute value of total bias (n = 1000). To measure the reduction of accuracy (and reward) caused by the total bias, we calculated the mean difference of accuracy in trials with low and high total bias in each session. Trials were labeled as low (or high) bias, if the absolute value of the total bias was within the smallest (or largest) tertiles of the distribution of absolute total bias across all sessions.

### Neuronal Data Analysis

#### Firing Rate and Dimensionality Reduction

In the present study we focused on the activity of PAG units before stimulus appearance, the period of time when the effects of choice biases could be most easily detected as stimulus cannot yet affect neuronal responses (Shadlen and Newsome, 2001). For each recorded unit, *m*, the firing rate (the number of spikes per unit of time) at trial *i, r*_*m*_(*i*), was computed in an 800 ms window that terminated 10 ms before dots onset.

Because of the large number of recorded units and limited number of trials, models that use neural responses to predict bias or choice are prone to overfitting. To diminish overfitting, we reduced dimensionality of the neuronal population activity using principal component analysis (PCA). The projection of the neural responses on the *j*^*th*^ principal component in the *i*^*th*^ trial is defined as

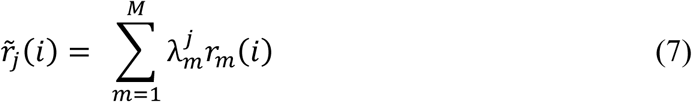

where 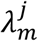 is the *j*^*th*^ PCA coefficient for the *m*^th^ unit and *M* is the number of simultaneously recorded units. For each recording session, we used the lowest number of PCA components, denoted *J*, which explained at least 50% of the total variance (range 38–52, median 48 components). The 50% cutoff provided a good balance between reducing overfitting (increasing prediction accuracy) and maintaining task-related variance of neural responses. Qualitatively similar results were obtained for different variance cutoffs or for the raw data. Principal components were calculated across all trials.

#### Decoding Biases — Linear Regression Models

A linear regression model was used to investigate whether PAG population activity represents fast, slow and total biases. We regressed any of these biases with the first *J* principal components of population activity as

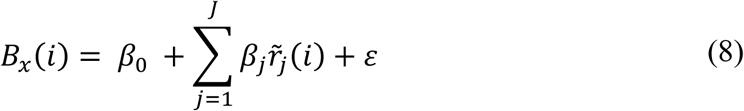

where “x” stands for {s, f, t} for slow (*B*_*s*_), fast (*B*_*f*_), or total bias (*B*_*t*_), respectively, and *ε* is a Gaussian noise term. *J* corresponds to the number of principal components that explain 50% of the total neural response variance, as explained above. Bias variables were z-scored using all trials in the session. The model was cross-validated using a leave-one-out procedure. The significance of the cross-validated *R*^*2*^ was assessed based on a permutation test (*n* = 1000 random shuffles of *B*_*x*_; the one-tailed *p*-value was computed as the fraction of shuffles leading to a *R*^*2*^ higher than the one obtained from unshuffled *B*_*x*_).

### Decoding Choices and Outcomes - Logistic Regression Models

We used a logistic regression model to predict the monkey’s *n*^*th*^ past choice, future choice, or outcome, based on population activity of the units before stimulus onset on the current trial.

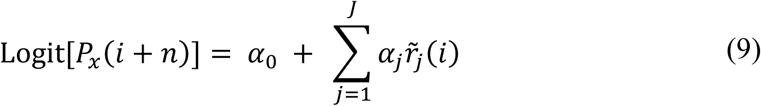

where *i* is the current trial, *n* ∈ {0, ±1, ±2, −3, −4, −5} and “x” stands for T1 for choice prediction and ‘+’ for outcome prediction (reward or not). Positive and negative n indicate trials after and before *i*, respectively.

For the choice decoder, we balanced the training set for T1 and T2 choices to make chance level equal to 0.5 for the model prediction accuracy. Similar to the behavioral model in Equation 4, for each repetition/cross-validation, we randomly removed trials corresponding to the surplus choice to have equal numbers of T1 and T2 choices. For the outcome decoder, we did not balance rewarded and unrewarded responses because errors comprised only a small fraction of trials (16%–29% across sessions) and balancing the number of rewarded and unrewarded trials led to exclusion of more than half of the trials in each session. Rather than dropping trials, we calculated the chance level for predicting trial outcome as the monkey’s overall reward rate (the fraction of the rewarded trials). Both models (choices or outcomes) were cross-validated using a leave-one-out procedure. For the model predicting upcoming choice (*n* = 0) the significance of the model prediction accuracy was assessed based on a permutation test similar to the ones described earlier (*n* = 1000 random shuffles of choices or outcomes).

For the remaining models we tested if the mean prediction accuracy calculated across sessions significantly differ from the chance level.

### Alignment of choice and the total bias decoders

The bias and choice decoders (Equations 8 and 9) provide distances of pre-stimulus responses from the discriminant hyperplanes that best explain the monkey’s choice or total bias (*d*_*choice*_and *d*_*bias*_). If the pre-stimulus choice and bias representations were aligned, predicting the upcoming choice based on *d*_*bias*_ would be as accurate as using both *d*_*choice*_and *d*_*bias*_. That is, *d*_*choice*_would not provide additional information for predicting the choice beyond what is provided by *d*_*bias*_.To test this hypothesis we trained and compared two logistic regression models:

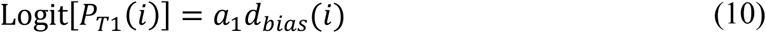

and

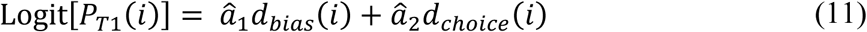

Alternatively, one could test for the alignment of the neural representations of choice and bias by directly comparing the two discriminant hyperplanes. If our definition of total bias and its neural representation provide a complete account of the representation of choice prior to stimulus onset, one would expect parallel hyperplanes in Equations 8 and 9, and thereby strong correlations in the weight vectors that determine the norm of hyperplanes (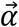 and 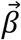). In contrast, if our definition of total bias is an incomplete account of the factors that predict the choice prior to stimulus onset, and if those factors have distinct neural representations from our total bias, the choice hyperplane would not align with our total bias hyperplane. Specifically, if we assume that factors beyond our total bias add up to make a new bias term, *ϑ*, that has a neural representation captured by 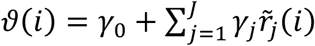, we can update Equation 8 to include all biases

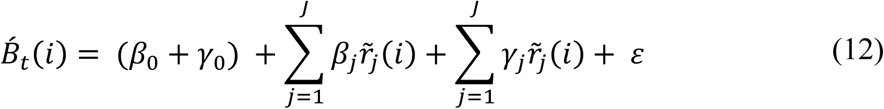

where 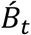 is the corrected bias term that includes both our slow and fast biases and additional factors that we may have failed to identify behaviorally in this paper. Because the combination of all possible bias terms is what enables the prediction of the upcoming choice based on neural responses prior to stimulus onset 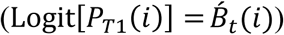, we can combine Equations 9 and 12 to write

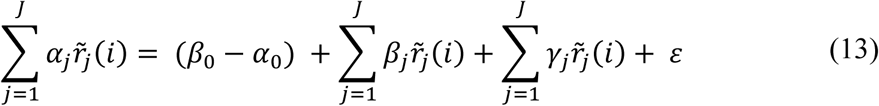

Equation 13 clarifies that in the presence of additional bias factors not captured by our definition of total bias, the choice hyperplane would not need to align well with our total bias hyperplane. Alternatively, if our definition of total bias is complete within the precision conferred by our dataset, the third term on the right-hand side of Equation 13 would be negligible and the choice and total bias hyperplanes would align well. We test for the alignment of hyperplanes by calculating the correlation of the weight vectors that determine their norms (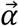 and 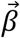), and by calculating the angle of the two vectors. Since both hyperplanes lay in a highly dimensional space, angles calculated are from two random hyperplanes are biased towards 90 deg (that is, they are biased to be orthogonal). To account for this bias, we also computed the angle between the two hyperplanes using only from two up to 38 top PCA dimensions used for the previous analysis. Additionally, we checked choice prediction accuracy and total bias representation in a reduced space.

In addition to the analyses above, we compared the accuracy of predicting choices using the neural representation of our total bias or the neural representation of choice prior to stimulus onset. For simplicity, we relied on the sign of *d*_*choice*_and *d*_*bias*_. Based on Equation 9, positive values of *d*_*choice*_mean a higher probability of choosing T1, while negative values mean a higher probability of choosing T2. Similarly, positive and negative *d*_*bias*_ suggest leaning toward T1 and T2, respectively.

### Choice decoder during stimulus presentation

To explore the dynamics of choice prediction accuracy based on neural responses, we extended Equation 9 to include time:

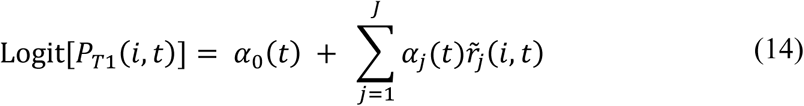

where *i* is the current trial and *t* is the center of time window used for the analysis. We used a 100-ms sliding window that moved from 800 ms before to 1100 ms after stimulus onset in steps of 20ms. The projection of the neural responses on the *j*^*th*^ principal component in the *i*^*th*^ trial 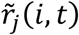 was defined using the same PCA coefficients used in Equation 7 and based on the 800 ms activity prior to stimulus onset. Similar to the model in Equation 9, we used a leave-one-out cross-validation and balanced choices in the training sets.

The model decision variable (DV) given by the right-hand side of the above equation is the distance of population neural responses from a linear discriminant hyperplane separating T1 and T2 choices. To investigate how behavioral bias interacts with the DV, we calculated the mean DV conditional on trials where the pre-stimulus total bias favored the choice finally made by the animal (matched trials) or on trials where the total bias was against the final choice (non-matched). To plot the average of DV for each group of trials, we flipped the sign of the DV for T2 choice trials before averaging.

## Supplementary Information

### Supplementary results

#### Impact of slow and fast biases on monkey’s performance

Since our task was designed such that the stimulus sequence across trials did not have temporal correlations, the existence of behavioral biases described above can only impair the monkey’s performance. As expected, periods with larger than average total bias correlated with periods with lower than average animal accuracy (percent of correct choices; Pearson correlation coefficient −0.08 ± 0.03, permutation test, *p*-value = 0.001). However, the reduction in accuracy was very small (0.007 ± 0.003%; one sample *t*-test, *p*-value = 0.02), suggesting that monkeys may not have noticed the adverse effect of biases on their performance or they did not find enough incentive to fully abolish them.

#### Analysis of angle between choice and total bias decoders

Conversion of the correlation coefficients to angles resulted in 70 ± 2.5 deg., significantly smaller than 90 (mean ± s.e.m., permutation test *p*-value = 0.001). However, we note that the angles were also far from zero, indicating that the alignment of the two hyperplanes was not perfect. The non-zero angles were at least partly due to the bias toward orthogonality in high-dimensional spaces. Repeating the correlation analysis and measuring angles in lower dimensional spaces (lower number of principal components) resulted in smaller angles as illustrated and elaborated in Supplementary Fig. 4. In any case, as demonstrated by our earlier analyses on choice prediction accuracy, the component of the choice hyperplane that was orthogonal to the bias hyperplane in the high-dimensional state space did not provide additional information for predicting behavior. Put differently, the non-zero angles between hyperplanes did not amount to functional differences with respect to the upcoming choice.

## Supplementary Figures

**Figure S1.**
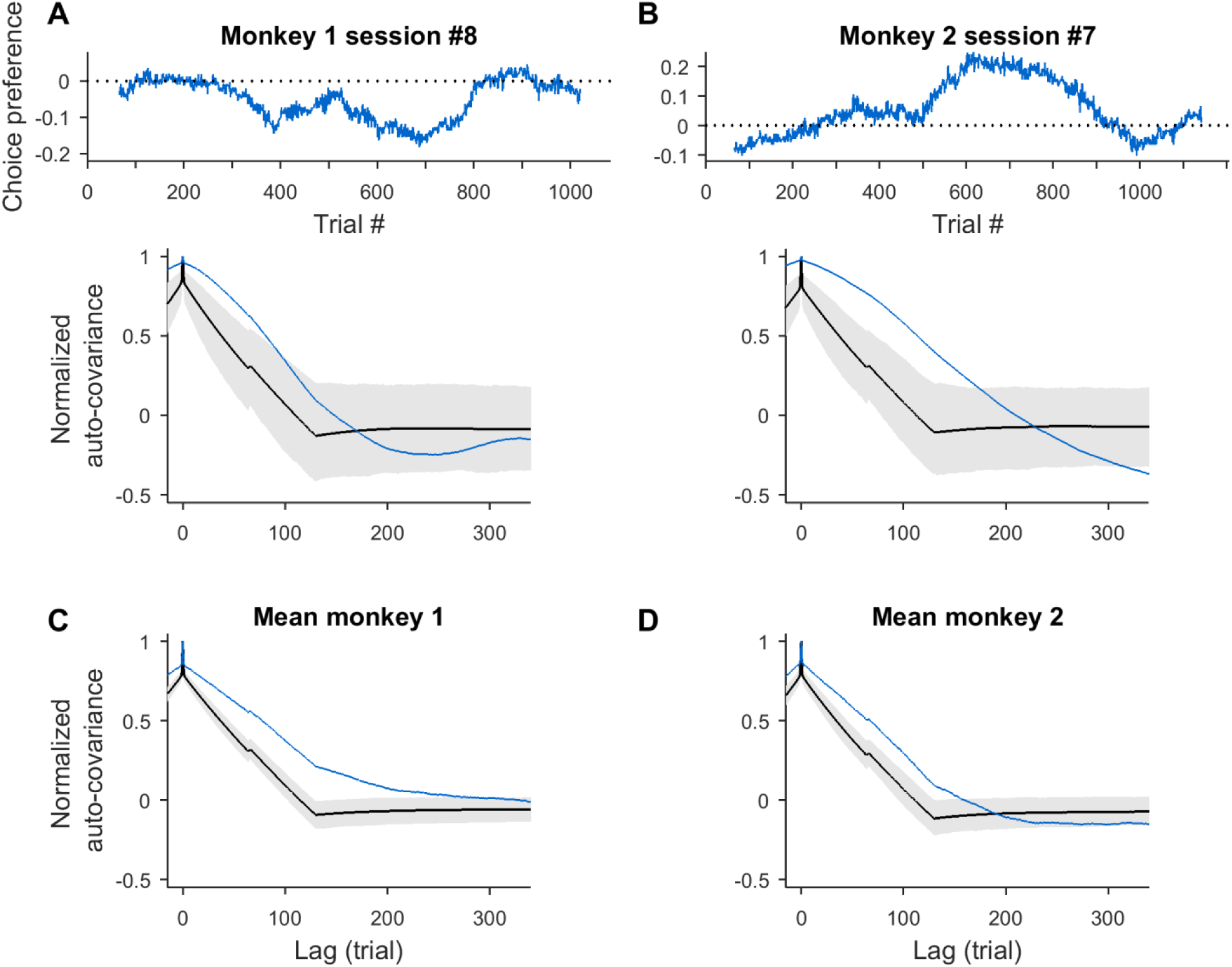
Statistically significant slow choice fluctuations through the session. **A – B** Slow fluctuations of choice preference are not an artifact generated by smoothing, as shown by the difference between the observed and shuffled-data auto-correlations. Top: slow choice preference calculated in 130 trials window (blue) from one example session of monkey 1 (A) and monkey 2 (B). The dotted line corresponds to condition with no choice preference. Bottom: the auto-correlograms of the slow choice preference fluctuation for experimental (blue) and shuffled data (black). The shuffled data were generated by permuting monkey choices across trials and recalculating the slow choice preference fluctuation (n = 1000). They represent the null hypothesis distribution when there are no actual fluctuations of choice preference. Black line corresponds to mean auto-correlograms of shuffled data and grey area corresponds to 90% of the distribution estimated from shuffles. C – D The mean auto-correlogram calculated across session for monkey 1 (C) and 2 (D). Convention as in A - B.

**Figure S2.**
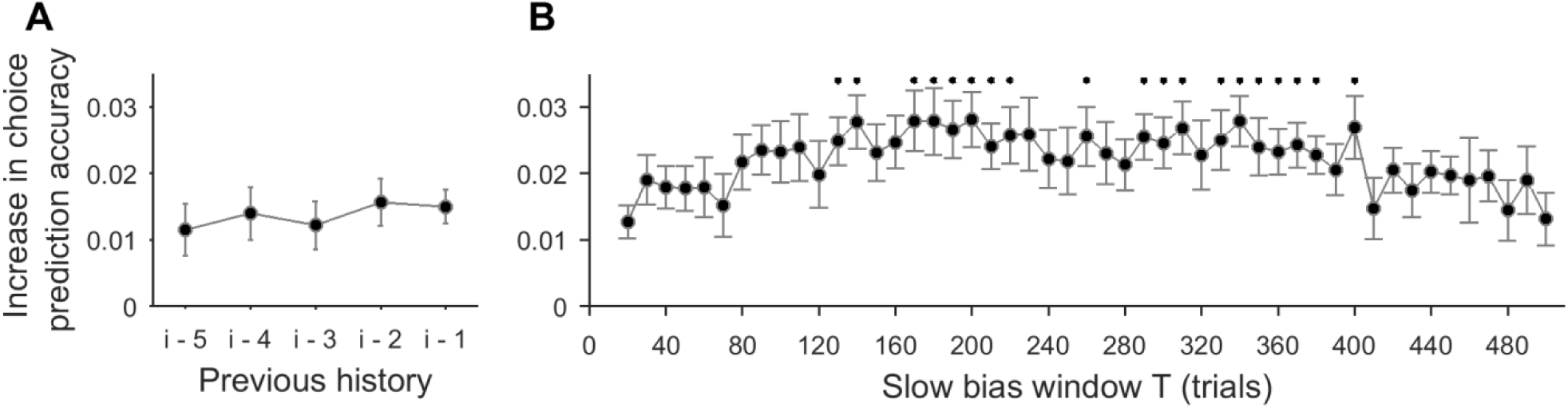
Slow and fast choice preference fluctuations have separated time scales. **A**. Mean increase in choice prediction accuracy of a model fitted to predict upcoming choice from the current stimulus coherence and the history of previous outcomes and choices in comparison to a model with just the coherence regressor. Fast choice preference is defined as a categorical variable that combines the choice and outcome in the previous trial but not on further trials (see Methods). Adding additional regressors from two trials back or further (i-k refers to a model including the coherence of the current trial i and choices and outcomes from the k previous trials as regressors) does not significantly improve the performance of the (i-1) model. **B**. The slow choice preference fluctuation occurs around a time scale of hundreds of trials. We took model (i-1) described before and extended it with one additional regressor - the choice preference calculated in a window T, varying between 20 and 500 trials. Significant improvement in the choice prediction accuracy of the extended model was observed for a broad range of T (*, 130 – 400 trials), but not outside this range. Significance tested using paired *t*-test (prediction accuracy for the (i-1) model against the extended model with slow choice preference).A-B. Prediction accuracy tested on difficult trials only.

**Figure S3.**
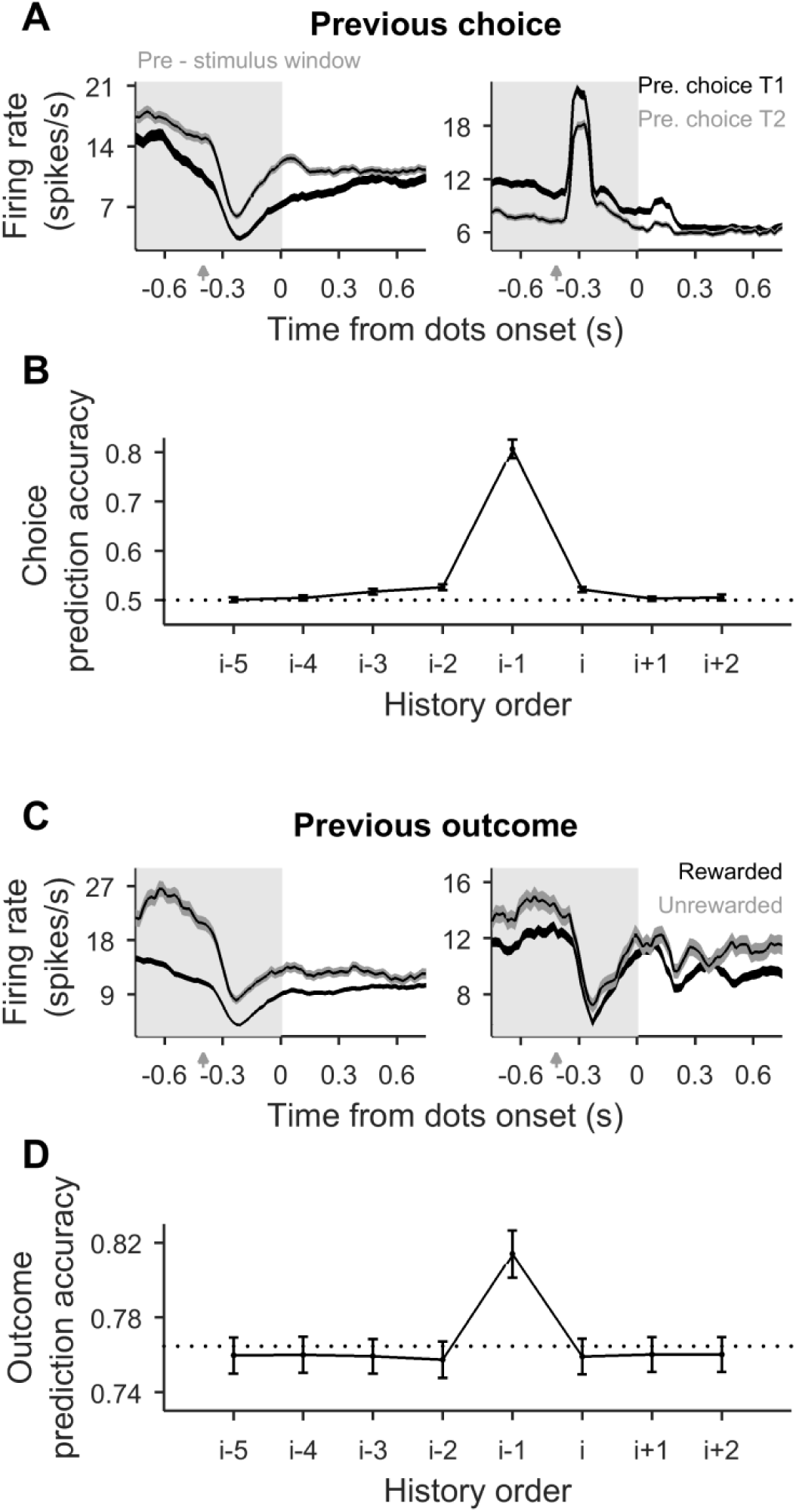
Pre-stimulus activity of PAG neurons caries information about previous choice and outcome – the two components of the fast behavioral bias. **A**. Firing rate of example cells (left - monkey 1 and right - monkey 2) averaged across trials when animal previously has chosen first (black) or second (gray) target. Shaded area around curves corresponds to SEM. Time zero refers to the onset of stimulus in the present (i^th^) trial. Spikes are counted in a 100 ms window swept with 20 ms resolution. **B**. Mean cross-validated prediction accuracy of decoding past and future choices from pre-stimulus activity of prearcuate gyrus cells. History and future horizons considered are up to five trials back and two trials forth from current i^th^ trial respectively. Rate calculated in 800 ms window before stimulus onset (shaded rectangle in A). Dotted black line marks chance level (0.5). Accuracy averaged across 16 sessions (± SEM). **C**. Firing rate of example cells averaged across previously rewarded (black) or unrewarded (gray) trials. **D**. Mean cross-validated prediction accuracy of decoding past and future outcome from pre-stimulus activity of prearcuate gyrus cells. Dotted black line marks chance level corresponding to animals performance (0.76, mean fraction of rewarded trials across 16 sessions). C – D convention like in A – B.

**Figure S4.**
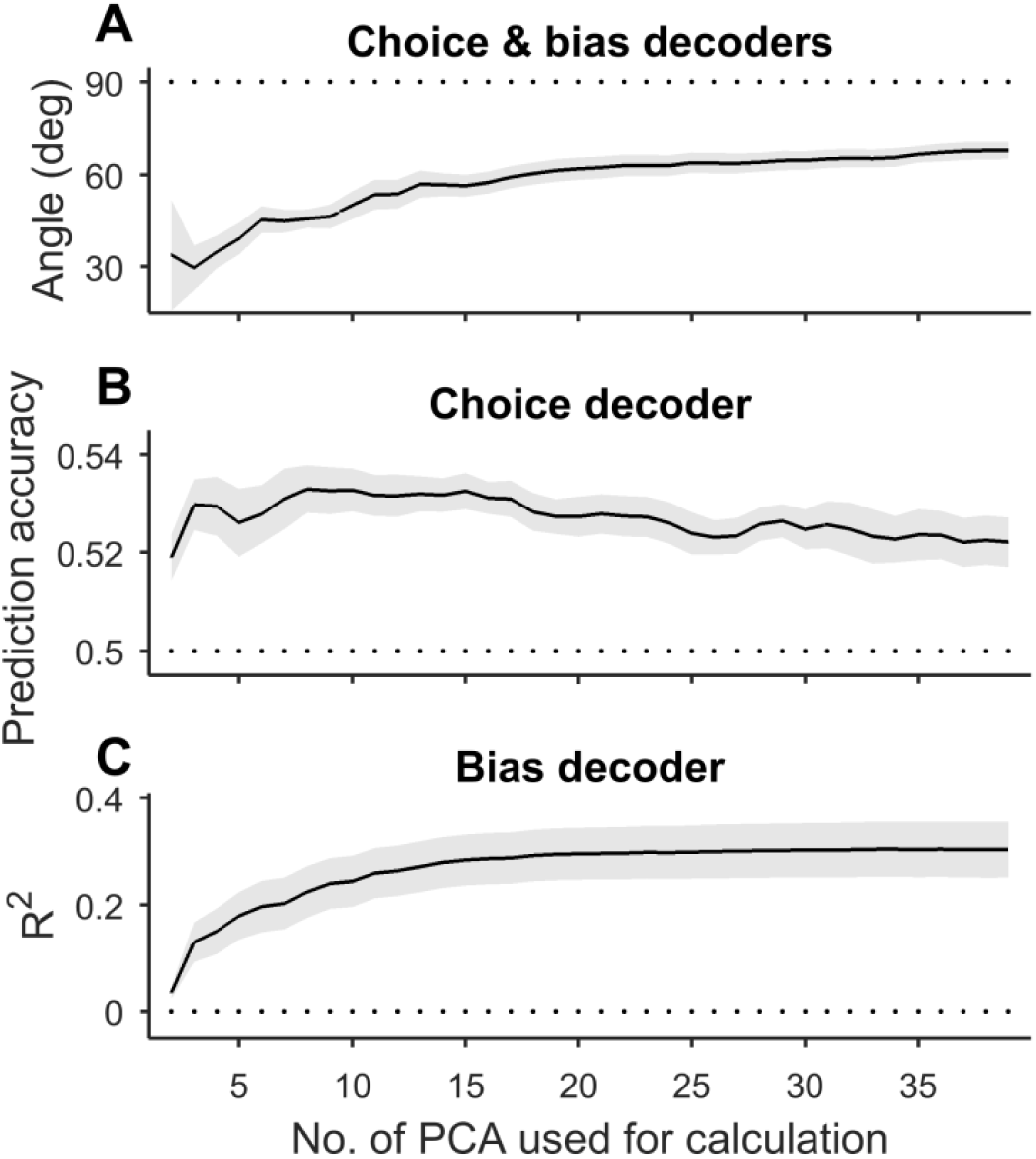
The statistics of choice and bias decoders as a function of number of principal components used in the analyses. **A**. Angle between weight vectors of choice and bias decoders **B**. Cross-validated prediction accuracy of choice decoder, and **C**. R^2^ of bias decoder calculated from test dataset of reduced dimensionality (from first two up to first 38 PCA components). Note that both decoders were trained on PCA’s explaining 50% of the variance.

## References

Abrahamyan, A., Silva, L.L., Dakin, S.C., Carandini, M., and Gardner, J.L. (2016). Adaptable history biases in human perceptual decisions. Proc. Natl. Acad. Sci. 113, E3548–E3557.

Akaishi, R., Umeda, K., Nagase, A., and Sakai, K. (2014). Autonomous Mechanism of Internal Choice Estimate Underlies Decision Inertia. Neuron 81, 195–206.

Akrami, A., Kopec, C.D., Diamond, M.E., and Brody, C.D. (2018). Posterior parietal cortex represents sensory history and mediates its effects on behaviour. Nature 554, 368–372.

Arandia-Romero, I., Nogueira, R., Mochol, G., and Moreno-Bote, R. (2017). What can neuronal populations tell us about cognition? Curr. Opin. Neurobiol. 46, 48–57.

Averbeck, B.B., Sohn, J.W., and Lee, D. (2006). Activity in prefrontal cortex during dynamic selection of action sequences. Nat Neurosci 9, 276–282.

Bogacz, R., Brown, E., Moehlis, J., Holmes, P., and Cohen, J.D. (2006). The physics of optimal decision making: A formal analysis of models of performance in two-alternative forced-choice tasks. Psychol. Rev. 113, 700–765.

Bonaiuto, J.J., Berker, A. de, and Bestmann, S. (2016). Changes in attractor dynamics predict altered perceptual decision making with dorsolateral prefrontal tDCS. BioRxiv 036905.

Britten, K.H., Shadlen, M.N., Newsome, W.T., and Movshon, J.A. (1992). The analysis of visual motion: a comparison of neuronal and psychophysical performance. J Neurosci 12, 4745–4765.

Busse, L., Ayaz, A., Dhruv, N.T., Katzner, S., Saleem, A.B., Schölvinck, M.L., Zaharia, A.D., and Carandini, M. (2011). The Detection of Visual Contrast in the Behaving Mouse. J. Neurosci. 31, 11351–11361.

Drugowitsch, J., and Pouget, A. (2012). Probabilistic vs. non-probabilistic approaches to the neurobiology of perceptual decision-making. Curr. Opin. Neurobiol. 22, 963–969.

Drugowitsch, J., Mendonça, A.G., Mainen, Z.F., and Pouget, A. (2019). Learning optimal decisions with confidence. Proc. Natl. Acad. Sci. 116, 24872–24880.

Eskandar, E.N., and Assad, J.A. (1999). Dissociation of visual, motor and predictive signals in parietal cortex during visual guidance. Nat. Neurosci. 2, 88–93.

Fan, Y., Gold, J.I., and Ding, L. (2018). Ongoing, rational calibration of reward-driven perceptual biases. ELife 7.

Fischer, J., and Whitney, D. (2014). Serial dependence in visual perception. Nat. Neurosci. 17, 738–743.

Fritsche, M., Mostert, P., and de Lange, F.P. (2017). Opposite Effects of Recent History on Perception and Decision. Curr. Biol. 27, 590–595.

Gardner, J.L. (2019). Optimality and heuristics in perceptual neuroscience. Nat. Neurosci. 22, 514–523.

Gigerenzer, G. (2008). Gut feelings: the intelligence of the unconscious (London: Penguin Books).

Gold, J.I., and Shadlen, M.N. (2007). The neural basis of decision making. Annu Rev Neurosci 30, 535–574.

Gold, J.I., Law, C.-T., Connolly, P., and Bennur, S. (2008a). The Relative Influences of Priors and Sensory Evidence on an Oculomotor Decision Variable During Perceptual Learning. J. Neurophysiol. 100, 2653–2668.

Gold, J.I., Law, C.-T., Connolly, P., and Bennur, S. (2008b). The Relative Influences of Priors and Sensory Evidence on an Oculomotor Decision Variable During Perceptual Learning. J. Neurophysiol. 100, 2653–2668.

Hall, P., Marron, J.S., and Neeman, A. (2005). Geometric representation of high dimension, low sample size data. J. R. Stat. Soc. Ser. B Stat. Methodol. 67, 427–444.

Hanks, T.D., Mazurek, M.E., Kiani, R., Hopp, E., and Shadlen, M.N. (2011). Elapsed Decision Time Affects the Weighting of Prior Probability in a Perceptual Decision Task. J. Neurosci. 31, 6339–6352.

Hastie, T., Tibshirani, R., and Friedman, J. (2001). The Elements of Statistical Learning (NY: Springer-Verlag).

Hayden, B.Y., and Moreno-Bote, R. (2018). A neuronal theory of sequential economic choice. Brain Neurosci. Adv. 2, 239821281876667.

Hermoso-Mendizabal, A., Hyafil, A., Rueda-Orozco, P.E., Jaramillo, S., Robbe, D., and Rocha, J. de la (2020). Response outcomes gate the impact of expectations on perceptual decisions. Nat. Commun. 11, 1–13.

Hesselmann, G., Kell, C.A., Eger, E., and Kleinschmidt, A. (2008). Spontaneous local variations in ongoing neural activity bias perceptual decisions. Proc. Natl. Acad. Sci. 105, 10984–10989.

Hwang, E.J., Dahlen, J.E., Mukundan, M., and Komiyama, T. (2017). History-based action selection bias in posterior parietal cortex. Nat. Commun. 8, 1–14.

Jasper, A.I., Tanabe, S., and Kohn, A. (2019). Predicting Perceptual Decisions Using Visual Cortical Population Responses and Choice History. J. Neurosci. 39, 6714–6727.

Kahneman, D. (2011). Thinking, fast and slow (New York: Farrar, Straus and Giroux).

Kiani, R., Hanks, T.D., and Shadlen, M.N. (2008). Bounded integration in parietal cortex underlies decisions even when viewing duration is dictated by the environment. J Neurosci 28, 3017–3029.

Kiani, R., Cueva, C.J., Reppas, J.B., and Newsome, W.T. (2014a). Dynamics of Neural Population Responses in Prefrontal Cortex Indicate Changes of Mind on Single Trials. Curr. Biol. 24, 1542–1547.

Kiani, R., Cueva, C.J., Reppas, J.B., and Newsome, W.T. (2014b). Dynamics of Neural Population Responses in Prefrontal Cortex Indicate Changes of Mind on Single Trials. Curr. Biol. 24, 1542–1547.

Kiani, R., Cueva, C.J., Reppas, J.B., Peixoto, D., Ryu, S.I., and Newsome, W.T. (2015). Natural Grouping of Neural Responses Reveals Spatially Segregated Clusters in Prearcuate Cortex. Neuron 85, 1359–1373.

Kim, J.-N., and Shadlen, M.N. (1999). Neural correlates of a decision in the dorsolateral prefrontal cortex of the macaque. Nat. Neurosci. 2, 176–185.

Link, S.W. (1992). The wave theory of difference and similarity (Hillsdale, NJ, US: Lawrence Erlbaum Associates, Inc).

Lueckmann, J.-M., Macke, J.H., and Nienborg, H. (2018). Can Serial Dependencies in Choices and Neural Activity Explain Choice Probabilities? J. Neurosci. 38, 3495–3506.

Mante, V., Sussillo, D., Shenoy, K.V., and Newsome, W.T. (2013). Context-dependent computation by recurrent dynamics in prefrontal cortex. Nature 503, 78–84.

Moreno-Bote, R., Shpiro, A., Rinzel, J., and Rubin, N. (2008). Bi-stable depth ordering of superimposed moving gratings. J Vis 8.

Moreno-Bote, R., Knill, D.C., and Pouget, A. (2011). Bayesian sampling in visual perception. Proc Natl Acad Sci U A 108, 12491–12496.

Moreno-Bote, R., Beck, J., Kanitscheider, I., Pitkow, X., Latham, P., and Pouget, A. (2014). Information-limiting correlations. Nat. Neurosci. 17, 1410–1417.

Nogueira, R., Abolafia, J.M., Drugowitsch, J., Balaguer-Ballester, E., Sanchez-Vives, M.V., and Moreno-Bote, R. (2017). Lateral orbitofrontal cortex anticipates choices and integrates prior with current information. Nat. Commun. 8, 1–13.

Padoa-Schioppa, C. (2013). Neuronal Origins of Choice Variability in Economic Decisions. Neuron 80, 1322–1336.

Peixoto, D., Kiani, R., Chandrasekaran, C., Ryu, S.I., Shenoy, K.V., and Newsome, W.T. (2018). Population dynamics of choice representation in dorsal premotor and primary motor cortex. BioRxiv 283960.

Rao, V., DeAngelis, G.C., and Snyder, L.H. (2012). Neural Correlates of Prior Expectations of Motion in the Lateral Intraparietal and Middle Temporal Areas. J. Neurosci. 32, 10063–10074.

Ratcliff, R., and Smith, P.L. (2004). A Comparison of Sequential Sampling Models for Two-Choice Reaction Time. Psychol. Rev. 111, 333–367.

Rustichini, A., and Padoa-Schioppa, C. (2015). A neuro-computational model of economic decisions. J. Neurophysiol. 114, 1382–1398.

Shadlen, M.N., and Kiani, R. (2013). Decision Making as a Window on Cognition. Neuron 80, 791–806.

Shadlen, M.N., and Newsome, W.T. (2001). Neural Basis of a Perceptual Decision in the Parietal Cortex (Area LIP) of the Rhesus Monkey. J. Neurophysiol. 86, 1916–1936.

Sohn, H., Narain, D., Meirhaeghe, N., and Jazayeri, M. (2019). Bayesian Computation through Cortical Latent Dynamics. Neuron 103, 934-947.e5.

Summerfield, C., and de Lange, F.P. (2014). Expectation in perceptual decision making: neural and computational mechanisms. Nat. Rev. Neurosci. 15, 745–756.

Urai, A.E., de Gee, J.W., Tsetsos, K., and Donner, T.H. (2019). Choice history biases subsequent evidence accumulation. ELife 8, e46331.

Williams, Z.M., Elfar, J.C., Eskandar, E.N., Toth, L.J., and Assad, J.A. (2003). Parietal activity and the perceived direction of ambiguous apparent motion. Nat. Neurosci. 6, 616–623.

Wyart, V., and Tallon-Baudry, C. (2009). How Ongoing Fluctuations in Human Visual Cortex Predict Perceptual Awareness: Baseline Shift versus Decision Bias. J. Neurosci. 29, 8715–8725.

Yu, B.M., Cunningham, J.P., Santhanam, G., Ryu, S.I., Shenoy, K.V., and Sahani, M. (2009). Gaussian-Process Factor Analysis for Low-Dimensional Single-Trial Analysis of Neural Population Activity. J. Neurophysiol. 102, 614–635.

